# The sensorimotor strategies and neuronal representations of tactile shape discrimination in mice

**DOI:** 10.1101/2020.06.16.126631

**Authors:** Chris C Rodgers, Ramon Nogueira, B Christina Pil, Esther A Greeman, Stefano Fusi, Randy M Bruno

## Abstract

Humans and other animals can identify objects by active touch, requiring the coordination of exploratory motion and tactile sensation. The brain integrates movements with the resulting tactile signals to form a holistic representation of object identity. We developed a shape discrimination task that challenged head-fixed mice to discriminate concave from convex shapes. Behavioral decoding revealed that mice did this by comparing contacts across whiskers. In contrast, mice performing a shape detection task simply summed up contacts over whiskers. We recorded populations of neurons in the barrel cortex, which processes whisker input, to identify how it encoded the corresponding sensorimotor variables. Neurons across the cortical layers encoded touch, whisker motion, and task-related signals. Sensory representations were task-specific: during shape discrimination, neurons responded most robustly to behaviorally relevant whiskers, overriding somatotopy. We suggest a similar dynamic modulation may underlie object recognition in other brain areas and species.

## Introduction

Animals have evolved sophisticated abilities to recognize objects, such as landmarks around food sources. Peripheral sensory neurons detect low-level object features and the central nervous system integrates them into a holistic representation of shape, endowed with behavioral meaning. This integration can be over time, space, or even multiple senses. Moreover, animals choose how to move their sensory organs to most effectively gather information about the world (Gibson, 1962; Yang et al., 2016c). A key challenge in neuroscience is to understand the strategies animals use to explore the world and how they integrate those motor actions with the resulting sensory input.

We investigated this problem in the mouse whisker system. Rodents rely on their whiskers (macrovibrissae) for social interaction and guiding locomotion (Grant et al., 2018; Gustafson and Felbain-Keramidas, 1977; Stüttgen and Schwarz, 2018). The whiskers, like human fingertips, are moved together onto objects in order to identify them (Ahissar and Assa, 2016; Diamond, 2010). Head-fixation permits precise quantification of whisker motion and contacts in high-speed video, as well as a wealth of modern techniques for monitoring and manipulating neural activity (Adesnik and Naka, 2018). Moreover, the individual columns of barrel cortex that process input from each whisker are readily identifiable *in vivo*. Thus, the whisker system is well-suited to the study of active touch, given an appropriate behavioral task.

The mouse’s ability to recognize novel objects is a model of cognition in health and disease (Lyon et al., 2012), but the underlying sensorimotor strategies and neuronal mechanisms are not understood. This is in part due to a paucity of suitable behavioral paradigms. On the one hand, freely moving rodents use their whiskers to identify objects and obstacles (Brecht et al., 1997; Hutson and Masterton, 1986; Voigts et al., 2015), but tracking multiple whiskers in freely moving animals is challenging (Petersen et al., 2020; Voigts et al., 2008). It is also difficult to ensure that freely moving rodents use only their whiskers, instead of vision, olfaction, or touch with skin (Mehta et al., 2007). On the other hand, most tasks for head-fixed mice focus on spatially simple features, like the location of a pole or the texture of sandpaper (Chen et al., 2013; O’Connor et al., 2010a). Indeed, the head-fixed mouse is often trimmed to a single whisker, though a few studies have considered multi-whisker behaviors (Brown et al., 2020; Celikel and Sakmann, 2007; Knutsen et al., 2006; Pluta et al., 2017).

We asked how mice discriminate objects of different curvature (concave or convex). Curvature is one of the fundamental components of form, and discriminating curvature requires integrating information over space (Connor et al., 2007; Lederman and Klatzky, 1987). Shape discrimination has never been studied with precise whisker tracking (although *cf*. Anjum et al., 2006; Brecht et al., 1997; Diamond et al., 2008; Harvey et al., 2001; Polley et al., 2005). Curved stimuli have been used in the visual and somatosensory systems of primates, but typically in passive presentation (Nandy et al., 2013; Yau et al., 2009). Active sensation is critical for shape discrimination in humans and other species (Chapman and Ageranioti-Bélanger, 1991; von der Emde, 2010; Klatzky and Lederman, 2011) yet the underlying neural mechanisms remain unknown.

We set out to understand the sensorimotor strategies and neuronal representations of shape discrimination. We trained head-fixed mice to discriminate concave and convex shapes while tracking every contact they made on the shapes with an array of whiskers. For comparison, other mice were trained simply to report the presence or absence of the same shapes regardless of their identity. We used behavioral decoding to reveal which sensorimotor features were critical for their decisions. Shape discrimination mice compared contacts across whiskers whereas shape detection mice simply summed up contacts across whiskers. Population recordings in barrel cortex revealed a persistent representation of the mouse’s choice on individual trials in addition to other sensory, motor, and task variables. Most importantly, during detection the neural population encoded contact by each whisker equally whereas during discrimination responses to behaviorally relevant whiskers were enhanced. Our multi-pronged approach of behavioral classification and neural encoding and decoding models reveals how the barrel cortex integrates fine-scale sensorimotor events into high-level representations of form.

## Results

### The shape discrimination and shape detection tasks

We developed a novel behavioral paradigm for head-fixed mice that challenged them to discriminate shapes (Supplemental Video 1). On each trial, a linear actuator moved a curved shape (either convex or concave) into the range of the whiskers on the right side of the face, though mice had to actively whisk to contact it. The shape stopped at one of three different distances from the mouse (termed close, medium, or far; Fig 1A,B), ensuring recognition was position-invariant and that mice did not memorize the location of a single point on the object. Mice could also generalize to flatter, more difficult stimuli (Supplemental Fig 1A).

**Figure 1.**
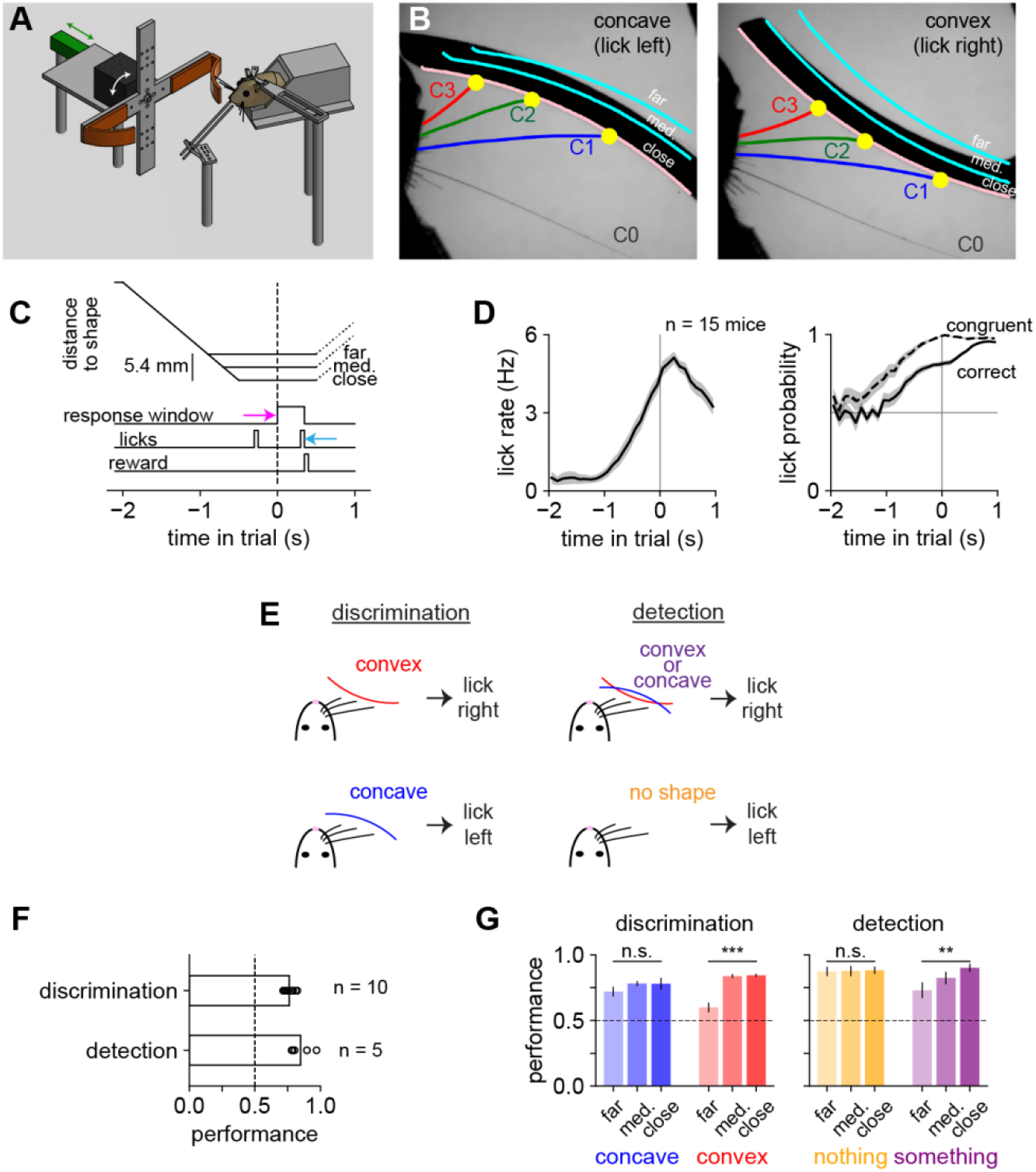
Mice learned either a shape discrimination or a shape detection task. **A)** Diagram of the behavioral apparatus. A motor (black) rotated a shape (orange) into position, and a linear actuator (green) moved it into the whisker field. Mice reported choice by licking left or right pipes. **B)** Example high-speed video frames. Shapes were presented at one of three different positions (pink and cyan lines labeled close, medium, and far). We tracked whiskers C0 (β or γ), C1, C2, and C3. **C)** Trial timeline. At t = −2 s, the linear actuator began moving the shape toward the mouse’s whisker field and reached its final position between t = −1.0 and −0.5 s. The response window opened at t = 0 s (pink arrow) and ended at the “choice lick” (cyan arrow). **D)** *Left*: the total lick rate (regardless of lick direction). *Right*: the probability that licks were correct (solid line) or congruent with the choice lick (dashed line). Correct licks increased with trial feedback (correct or incorrect) after t = 0 s. **E)** Task rules. Mice trained on shape discrimination licked right for convex and left for concave shapes. Mice trained on shape detection licked right for either shape and left on trials when no shape is presented. **F)** Mouse performance (fraction of correct trials) on both tasks exceeded chance (dashed line). **G)** Mouse performance by task, stimulus, and position. On the “nothing” condition, the actuator moves to the correct position, but no shape is present. Performance significantly improves with proximity on convex shapes during discrimination and on any shape during detection (one-way repeated-measures ANOVA). Throughout the manuscript: * p < 0.05; ** p < 0.01; *** p < 0.001. Error bars: SEM over mice.

Mice learned to lick left for concave and right for convex shapes in order to receive a water reward. Two seconds after the shape started moving, and soon after it reached its final position, the “response window” began (Fig 1C). The first lick in the response window (the “choice lick”) determined whether the trial was correct or incorrect. Early licks had no effect, but mice increased their rate of correct licks and licks concordant with the eventual choice lick as the response window approached (Fig 1D), indicating the formation of their decision. Mice could learn the trial timing from the sound of the actuator, but whiskers were required to identify the shape (Supplemental Fig 1B).

To unambiguously identify each whisker in videography, we gradually trimmed off whiskers on the right side of the face throughout training. The middle (C) row was spared: C1 is the caudal-most and longest whisker; C3 is the rostral-most and shortest whisker still capable of reaching the shapes. Mice were strongly impaired by each trim, falling to chance or near-chance levels, suggesting that they initially relied on many whiskers (data not shown). However, with retraining, many were able to discriminate shape with only these 3 whiskers. Some mice retained a straddler whisker (“C0”), but it rarely made contact and was excluded from analysis.

We trained a separate group of mice on a “shape detection” task (Fig 1E) to determine whether the behavioral and neural responses were specific to shape discrimination or were simply due to the shapes themselves. In this control task, mice learned to lick right in response to either shape and to lick left on trials when the actuator presented an empty position with no shape. The shapes, trial timing, and trimming were identical to those for discrimination.

Both groups of mice learned to perform well above chance (Fig 1F; n = 5 detection mice and 10 discrimination mice). Detection mice more accurately reported the presence of a shape when it was closer (Fig 1G). Discrimination mice identified concave shapes equally well at all locations, but were more likely to identify convex shapes correctly when closer. Thus, shape discrimination relied on “detecting convexity”, an observation we return to below.

### The whisk cycle synchronizes contacts across multiple whiskers into packets

To identify how mice identified the shapes, we acquired video of their whiskers at 200 frames per second. This large dataset—15 mice, 88.9 hours, 115 sessions, 18,514 trials, 63,979,800 frames—necessitated high-throughput automated tracking. To do this, we used the human-curated output of a previous-generation whisker tracking algorithm (Clack et al., 2012) to bootstrap the training of a deep convolutional neural network (Insafutdinov et al., 2016; Mathis et al., 2018; Pishchulin et al., 2015). This method successfully tracked the full extent of the whiskers even as they moved rapidly, became obscured, or contacted the shape (example frames: Supplemental Fig 2A).

Trained mice whisked in stereotyped patterns that could differ widely across individuals (Fig 2A). We decomposed whisker motion into individual cycles (Fig 2B, n = 882,893 whisks from 15 mice, excluding inter-trial intervals). Individual whisks had a mean duration of 64.1 ± 4.0 ms, equivalent to a whisking frequency of 15.6 Hz, with an amplitude (peak-to-trough angular difference) of 10.6 ± 1.9° (mean ± standard deviation of the within-mouse average; Supplemental Fig 2B). Mice made contacts near the peak of the whisk cycle (Fig 2C), synchronously across whiskers (Fig 2D; *cf.* Sachdev et al., 2001).

**Figure 2.**
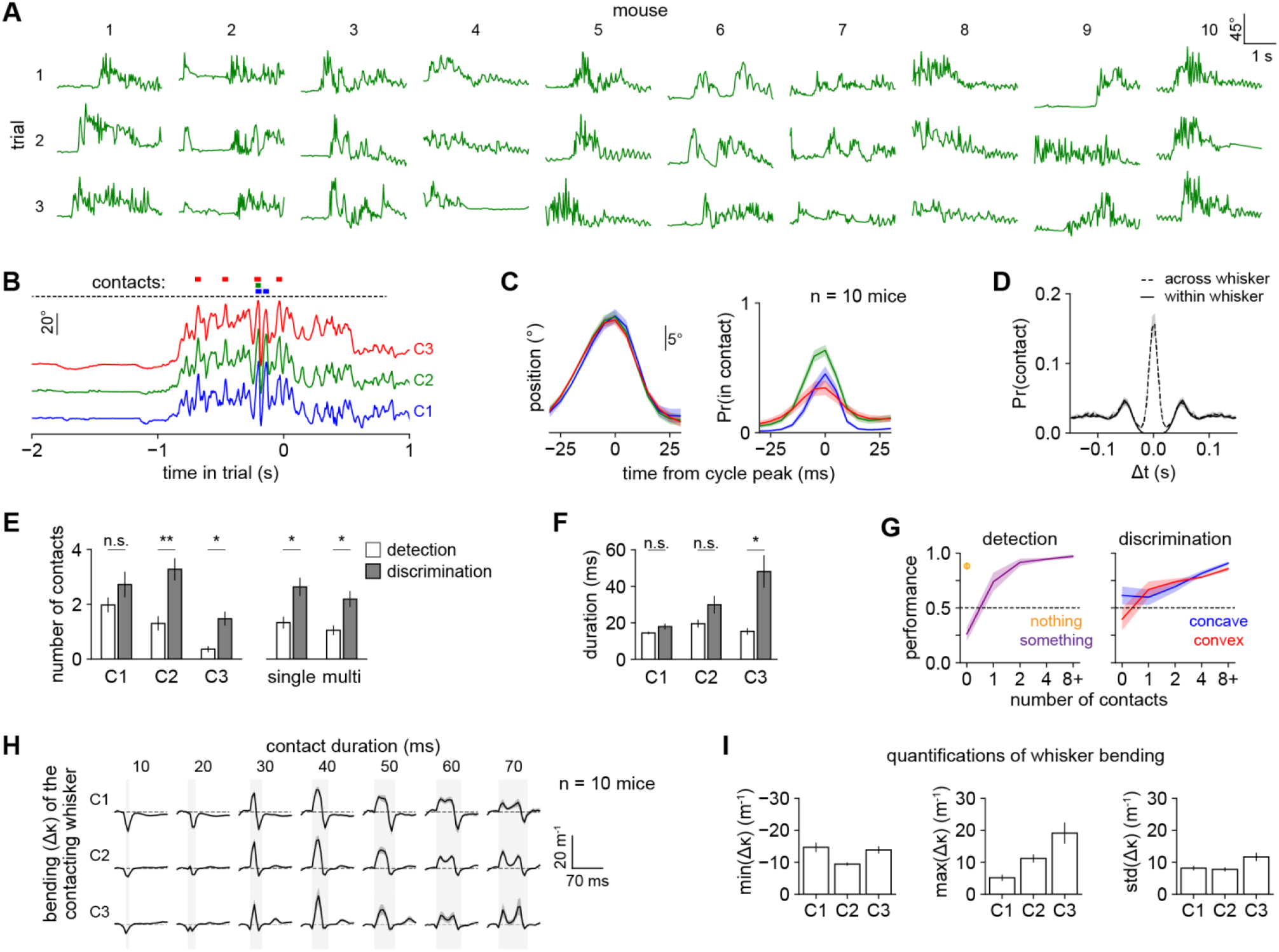
Mice employ brief taps with multiple whiskers to discriminate shape. **A)** Angular position of the C2 whisker on three representative correct trials from ten mice performing shape discrimination. **B)** Angular position of C1, C2, and C3 over a single trial using timescale defined in Fig 1C. Colored bars: whisker contacts. **C)** *Left*: mean angle of each whisker aligned to the C2 whisk cycle peak. *Right*: probability that each whisker was in contact, aligned to the same time axis as on left. For both, n = 94,999 whisk cycles during which ≥1 whisker made contact. **D)** Autocorrelation of contact times within each whisker (solid) and cross-correlation of contact times across pairs of adjacent whiskers (dashed), expressed as probability of contact at each lag. **E)** *Left*: the mean number of contacts made by each whisker during both tasks. Discrimination mice made significantly more contacts with C2 and C3 than detection mice did. *Right*: Discrimination mice made significantly more contacts with a single whisker alone and with multiple whiskers simultaneously than detection mice did (two-sample t-test). **F)** Mean duration of contacts in panel E. C3 contacts are significantly longer during discrimination (two-sample t-test). **G)** Performance versus the number of contacts in the detection (left) or discrimination (right) task. Multi-whisker contacts were counted as a single contact event. Orange circle: trials during detection when no shape is present. We excluded mice from any bin in which they had <10 trials. **H)** Mean whisker bending (Δκ) over time during each contact aligned to its onset and relative to the pre-contact baseline (dashed line), plotted separately for each whisker (row) and contact duration (column). Shaded area: duration of contact. Not all mice made contacts of all possible durations; data points with <10 contacts per mouse were excluded. **I)** Whisker bending quantified as the minimum, maximum, and standard deviation of Δκ over the duration of each contact. Error bars: SEM over mice. All panels include 10 discrimination mice. E-G also include 5 detection mice.

Compared with the detection group, mice performing shape discrimination made more single- and multi-whisker contacts (Fig 2E). Both groups made C3 contacts less frequently because it was too short to touch the shapes at the further positions. However, the shape discrimination group made much longer duration contacts with the C3 whisker than the shape detection group (Fig 2F), suggesting an important role for this whisker in discrimination. During both tasks, performance increased with the number of contacts made on each trial (Fig 2G). In combination with the stereotyped whisking pattern, this suggests mice relied on a pre-planned motor strategy rather than an closed-loop strategy (Yang et al., 2016c; Zuo and Diamond, 2019a). In sum, the whisk cycle synchronizes contacts across multiple whiskers into discrete packets of sensory evidence, which mice use to identify shape.

### Mice rely on brief “tapping” of the stimuli

The way mice contacted these shapes fundamentally differed from previous reports of mice and rats exploring different objects. We exclusively observed tip contact whereas mice localizing poles make contact with the whisker shaft (Hires et al., 2013, *cf.* a similar observation in rats discriminating texture in Carvell and Simons, 1990). We never observed mice dragging their whiskers across the objects’ surfaces, as they do with textured stimuli (Carvell and Simons, 1990; Jadhav et al., 2009; Ritt et al., 2008).

Contacts were brief (median 15 ms, IQR 10-25 ms, n = 167,217; Supplemental Fig 2C). Whisker bending, a commonly used proxy for contact force (Birdwell et al., 2007) but see also (Quist et al., 2014), was dynamic (Fig 2H): a whisker could bend slightly while pushing into a shape and then bend in the other direction while detaching. Occasionally we observed double pumps, a signature of active exploration (Wallach et al., 2020).

Strikingly, the contact forces we observed were much smaller than in previous reports of other tasks. The typical maximum bend (Δκ) was 5.1 +/− 1.0 m^−1^ for C1, 11.2 +/− 1.2 m^−1^ for C2, and 19.1 +/− 3.3 m^−1^ for C3 (mean +/− SEM over mice; Fig 2I), much less bent than the 50-150 m^−1^ typical of pole localization or detection (Hires et al., 2015; Hong et al., 2018; Huber et al., 2012). This sensorimotor strategy of “minimal impingement” onto the shape is the mode used by freely moving rodents investigating objects and may thus be more naturalistic (Grant et al., 2009; Mitchinson et al., 2007).

### Behavioral decoding reveals the sensorimotor features that guide behavior

To uncover the strategies mice used to perform this novel task, we turned to behavioral decoding. First, we quantified a large suite of sensorimotor features from the video (*e.g.*, contact location, cross-whisker contact timing) as well as task-related variables (choice and reward history). Then we trained linear classifiers using logistic regression to predict either the stimulus identity (concave vs convex for discrimination; something vs nothing for detection) or the mouse’s choice (lick left or lick right) on each trial using those features (Fig 3A).

**Figure 3.**
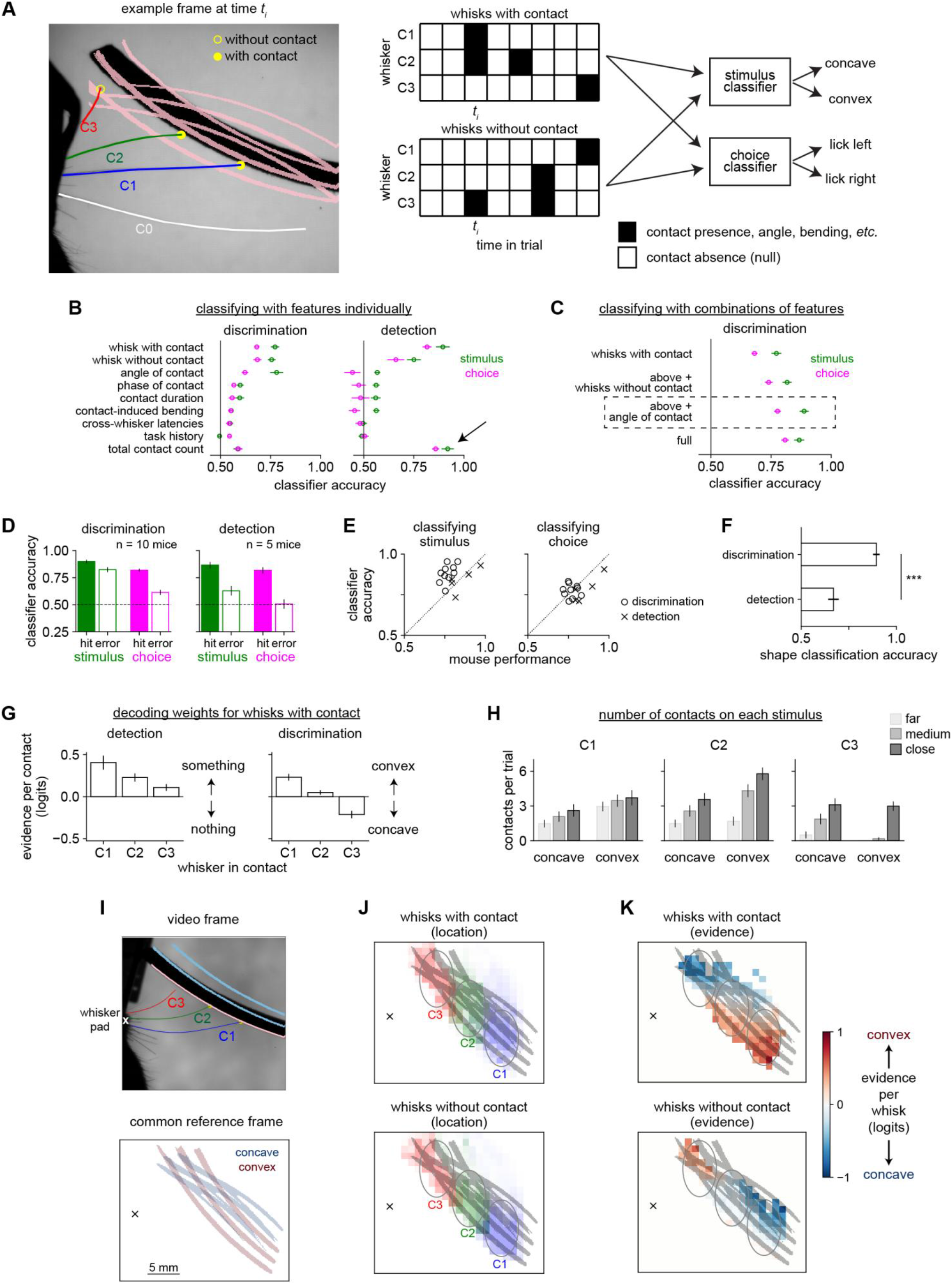
Mice compare contacts across whiskers in order to determine shape. **A)** A large suite of behavioral features was used to train behavioral decoders to predict the stimulus (concave or convex) or choice (direction of choice lick). *Left:* Example frame showing the peak of a whisk in which whiskers C1-C3 protracted far enough to reach the shapes at some positions (pink lines). In this example, C1 and C2 were scored as “with contact” and C3 as “without contact”. All three are considered “sampling whisks” because they protracted far enough to reach the shapes at their closest position. *Middle*: Features were extracted into sparse two-dimensional arrays of whisker (rows) versus 250 ms time bins (columns). Black squares indicate a whisk with contact (top) or without contact (bottom). Features could be binary (*e.g.*, contact by a specific whisker) or continuous (*e.g.*, angle or force). *Right*: Logistic regression classifiers predicted stimulus or choice. **B)** Feature importance quantified by the accuracy of a behavioral decoder trained on that feature alone to identify stimulus (green) or choice (pink). During shape detection (right), the total number of contacts (black arrow) was the most informative feature; this same parameter was much less useful during discrimination. **C)** Accuracy of decoders trained on combinations of features: first whisks with contact only, then also including whisks without contact, then including angle of contact, then including all features in the entire dataset. The third model (dashed box, “optimized behavioral decoder”) is used for the remainder of this figure, because it performs as well as the full model while using far fewer features. **D)** The optimized behavioral decoder predicts stimulus (green) and choice (pink) well during both shape discrimination and detection. Filled bars: correct trials; open bars: incorrect trials. **E)** Accuracy of the decoder (y-axis) versus performance of each mouse (x-axis) when decoding the stimulus (left) or choice (right) during discrimination (circles) or detection (X’s). The decoder identifies stimulus (left panel) significantly more accurately than the mouse does (left panel) during discrimination (p < 0.001, paired t-test) but not during detection (p > 0.05). **F)** Accuracy of decoders trained to distinguish concave from convex shapes using data from shape discrimination (top) or shape detection (bottom) tasks. The decoders were significantly more able to identify shape during discrimination (p < 0.001, unpaired t-test), indicating that the whisking strategy mice employed for shape discrimination extracted more information about the shape’s identity. **G)** The weights assigned by the decoder to the “whisks with contact” feature, separately plotted by which whisker made contact. Weights were relatively consistent over the trial timecourse (data not shown) and are averaged over time here for clarity. They are expressed as the change in log-odds (logits) per additional contact. **H)** The mean number of contacts per trial for each whisker during shape discrimination, separately by shape identity (concave or convex) and position (far, medium, or close indicated by shading; *cf.* Fig 1B). Although each whisker may touch one shape or the other more frequently, no whisker touches a single shape exclusively. **I)** Videos for all sessions and mice were registered into a common reference frame based on shape positions. *Top*: single frame showing whisker identity and location of whisker pad for reference. *Bottom*: location of the concave (blue) and convex (red) shapes in the common reference frame, with average location of whisker pad marked. **J)** Location of the peak of each whisk with contact (top) or without contact (bottom) in the common reference frame. Each whisker (C1, C2, and C3; blue, green, and red) samples distinct regions of shape space (gray ovals). **K)** The same data from panel (J), but now colored by their strength of the evidence about shape (red: convex; blue: concave) using the decoder weights. *Top:* C1 (C3) contacts occur in a region that is more likely to contain a convex (concave) shape. *Bottom:* On whisks without contact, the mapping between whisker and shape identity is reversed. Note that the sampled areas can contain either shape, so sampling one area alone is not sufficient to perform the task. Error bars: SEM over mice.

Predicting the stimulus indicated which features carried information about shape whereas predicting choice indicated which features might have influenced the mouse’s decision. However, an important challenge was to disentangle the extent to which each feature predicted stimulus or choice (Nogueira et al., 2017). These two variables are correlated; indeed, they are perfectly correlated on correct trials. To directly address this, we weighted error trials in inverse proportion to their abundance, such that correct and incorrect trials were balanced (*i.e.*, equally weighted in aggregate). This notably improved our ability to predict the mouse’s errors (Supplemental Fig 3A).

To identify the most important features, we compared the accuracy of separate decoders trained on every individual feature during shape discrimination (Fig 3B, left). The most informative feature for decoding both stimulus and choice was a two-dimensional binary array representing <u>which</u> whisker made contact at each timepoint within the trial, which we term “whisks with contact” (schematized in Fig 3A). The next most informative feature was “whisks without contact”: when the mouse whisked far enough forward to rule out the presence of some shapes but did not actually make contact. Together, these two variables constitute all “sampling whisks” that were sufficiently large to reach the closest possible shape position; the remaining “non-sampling whisks” could not be informative because they were too small to reach the shapes at any position. The “contact angle” feature was also useful for predicting the stimulus, likely due to the geometrical information it contains. It was less useful for predicting choice, suggesting that mice did not exploit the information despite its utility.

The remaining 28 analyzed features were relatively uninformative about choice (Supplemental Fig 3B). Notably, mechanical/kinematic variables like speed or contact-induced whisker bending, cross-whisker variables like relative timing, and task variables like choice history contained little or no information about stimulus and choice. Similarly, contact time within the trial or whisk cycle was relatively uninformative compared with the spatiotemporal “whisks with contact” feature.

We next assessed whether these features contained unique information and which features sufficed for maximal prediction accuracy by gradually adding features in decreasing order of their usefulness until the model’s performance plateaued (Fig 3C). Unique information was present in whisks with contact, whisks without contact, and contact angle, and these three features together performed as well as the full model with all measured features. Therefore we used this reduced model (the “optimized behavioral decoder”; dashed box, Fig 3C) for all further analyses.

This decoder accurately predicted either stimulus or choice on both correct and error trials during both detection (Fig 3D; stimulus: 84.0 ± 3.2%; choice: 78.3 ± 3.6%; mean ± SEM) and discrimination (stimulus: 88.2 ± 1.8%; choice: 77.4 ± 1.3%). It outperformed the mice on shape discrimination (Fig 3E), indicating that the mice were unable to access or use some of the information in the contact pattern.

Thus, this decoder constitutes a model of behavior capable of either identifying the stimulus or predicting the mouse’s choice, even on error trials. Reflecting the mice’s “tapping” strategy, this model primarily required binary information about which whiskers made contact rather than the fine temporal dynamics of those contacts.

### Shape detection and discrimination engage distinct motor strategies

The results from classifying stimulus and choice in the shape detection task differed strikingly from shape discrimination: the total contact count summed over whiskers explained stimulus and choice better than any other variable (Fig 3B, right). Total contact count was far less informative during discrimination. This reflects the fundamental difference between these tasks: detection requires the mouse only to know that contacts occurred whereas discrimination requires additional information—most critically, the identities of the contacting whiskers.

To test whether mice adapted their whisking strategies to the task, we asked whether shapes could be classified from the data of mice performing the shape detection task, even though these mice did not actually need to identify the shapes. We used the optimized behavioral decoder for discrimination (Fig 3C, dashed box) to predict shape identity from detection sessions. Its ability to decode shape identity during the detection task was poor compared with during the discrimination task (Fig 3F), despite the fact that the shapes in both cases were identical. Thus, mice adapt their whisking to the task at hand, collecting more information about shape identity when behaviorally relevant.

### Mice compare contacts across whiskers to discriminate shape

Whether predicting stimulus (Fig 3G) or choice (Supplemental Fig 3C), the behavioral decoder assigned strikingly different weights to contacts made by each whisker. For shape detection, all weights were positive, meaning contact by *any* whisker indicated that a shape is present (Fig 3G, left). In sharp contrast, weights of different whiskers had opposite signs during shape discrimination (Fig 3G, right). This indicates that they conveyed opposite information: each C1 contact indicated a greater likelihood of convex whereas each C3 contact indicated a greater likelihood of concave.

Thus, mice compare contacts across whiskers to discriminate an object’s curvature whereas they sum up contacts across whiskers to detect an object. Critically, this is not because any given whisker can only reach one of the shapes; all whiskers can touch both shapes (Fig 3H). Instead, the whisking strategy employed for discrimination biases contact prevalence across whiskers, which the decoder exploits to predict the mouse’s choice.

To visualize this process of spatial sampling, we registered all of our whisker video into a common reference frame (Fig 3I). As expected, the whiskers reliably sampled different regions of shape space (Fig 3J). Interestingly, the C1 whisker sampled the region in which contacts indicated convexity and absence of contacts indicates concavity (Fig 3K). The reverse was true for C3. The location that mice chose to sample even in the absence of contacts was also informative about their upcoming choice (Supplemental Fig 3D-F; Dominiak et al., 2019).

In summary, behavioral decoding produced a computational model of the distinct sensorimotor strategies that mice adopted in two different tasks. Inspection of the weights revealed that mice summed up contacts across whiskers to detect shapes whereas they compared contacts across whiskers to discriminate shape identity. This analysis could be used to dissect active sensation in other modalities as well.

### Barrel cortex neurons encode movement, contacts, and choice

We next examined how barrel cortex encoded these sensorimotor events as well as cognitive variables like reward history, using an extracellular electrode array to record across all layers of cortex simultaneously (Fig 4A-D; Supplemental Video 2). We recorded 675 neurons from 7 mice performing shape discrimination and 301 neurons from 4 mice performing shape detection. Putative inhibitory interneurons were identified from their narrow waveform width (Fig 4B). Neurons with broad waveforms are mostly excitatory, though a small population of inhibitory neurons are similarly broad (Bruno and Simons, 2002; Gouwens et al., 2019; Yu et al., 2019).

**Figure 4.**
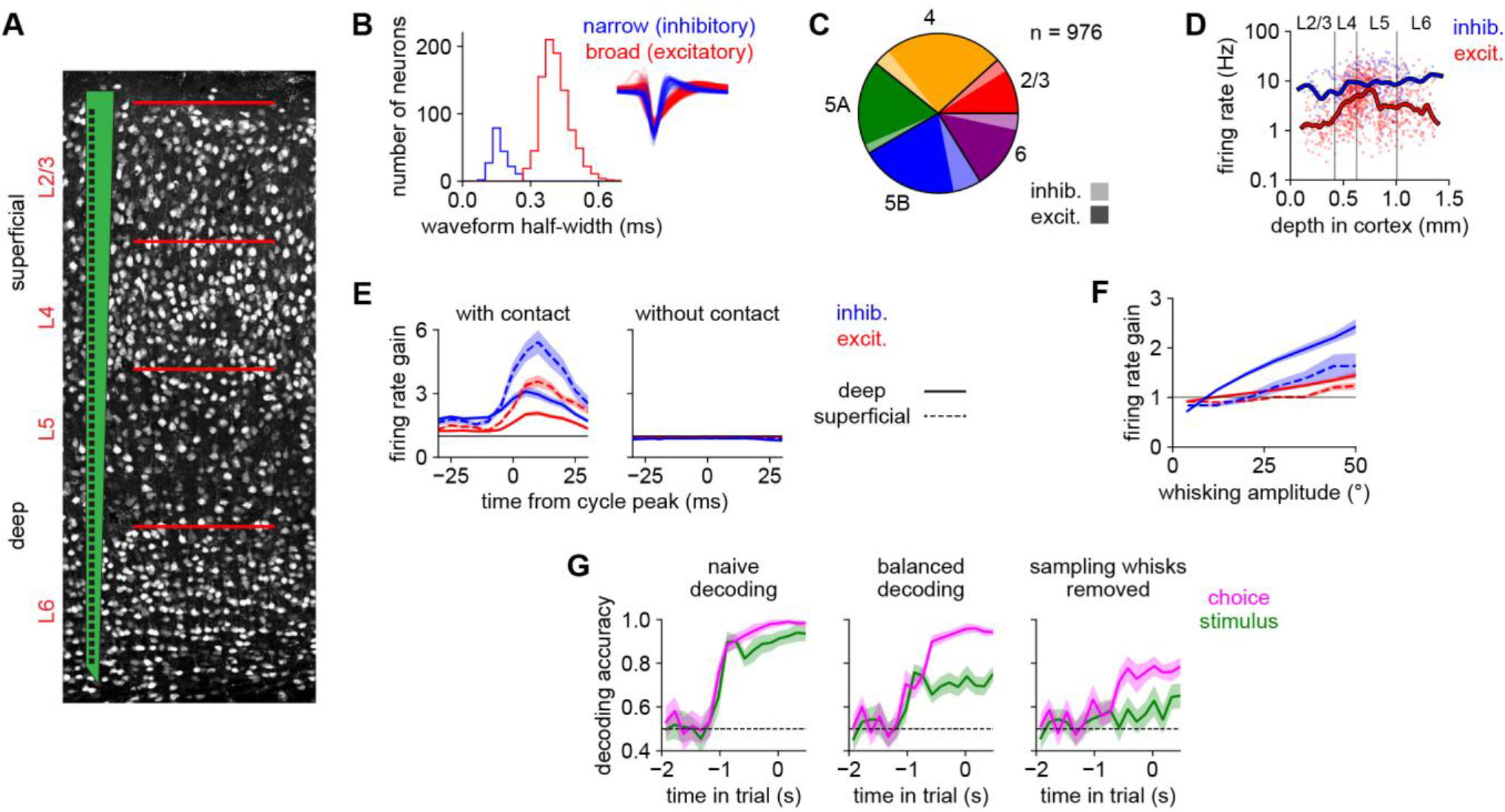
Barrel cortex neurons respond to whisking and contacts. **A)** Schematic of the multi-electrode array we used that spans all layers of cortex. Background: barrel cortex neurons labeled with NeuN. **B)** The distribution of extracellular waveform half-widths (*i.e.*, the time between peak negativity and return to baseline) of recorded neurons. The distribution is strongly bimodal, permitting nearly unambiguous classification into narrow-spiking (putative inhibitory; blue) and broad-spiking (putative excitatory; red) cell types. Inset: normalized average waveforms from individual neurons. In panels B-F, n = 976 neurons from both tasks, pooled because the results were similar. **C)** Relative fraction of excitatory (dark) and inhibitory (light) neurons recorded in each layer. **D)** Mean firing rates over the entire session of individual neurons versus their depth in cortex. Inhibitory and deep-layer neurons typically exhibit higher firing rates. Lines: smoothed with a Gaussian kernel. **E)** Firing rate gain (responses divided by each cell’s mean firing rate over a session) locked to the whisk cycle for superficial (L2/3 and L4; dashed lines) and deep (L5 and L6; solid lines) neurons, separately plotted for whisks with contact (left) and without contact (right). Absolute firing rates shown in Supplemental Fig 4. Error bars: SEM over neurons. **F)** Firing rate gain of each cell type on individual whisk cycles without contact (y-axis) versus the amplitude of that whisk cycle (x-axis). Deep inhibitory neurons (solid blue line) are modulated most strongly. Error bars: SEM over neurons. **G)** Stimulus (green) or choice (pink) can be decoded from a pseudopopulation (n = 450 neurons) aggregated across shape discrimination sessions (timescale defined in Fig 1C). *Left*: With a naive (unbalanced) approach, stimulus or choice can be decoded with similar accuracy. *Middle*: Equally balancing correct and incorrect trials decouples stimulus and choice. *Right*: Removing spike counts from all sampling whisks (*i.e.*, whisks sufficiently large to reach the shapes) largely abolishes stimulus information while choice information remains. Dashed line: chance. Error bars: 95% bootstrapped confidence intervals.

Neurons exhibited rapid transient responses to whisks with contact but not to whisks without contact (Fig 4E). These contact responses were stronger in the superficial layers and in inhibitory neurons, likely reflecting greater thalamocortical input to this cell type (Bruno and Simons, 2002; Cruikshank et al., 2007). Firing rates on timescales longer than the whisk cycle nevertheless tracked whisking amplitude, especially in deep inhibitory neurons (Fig 4F).

To determine whether these neurons contained information about stimulus or choice, we used neural decoding of resampled pseudopopulations to predict stimulus and choice from the entire neural population (Methods; Rigotti et al., 2013). Stimulus and choice could be decoded with a similar timecourse and accuracy (Fig 4G, left) but, as we addressed in the behavioral analysis, this could be an artifact of the correlation between them. We therefore applied a similar approach of “balancing”, *i.e.* weighting error trials in inverse proportion to their abundance, to completely decorrelate stimulus and choice. This revealed that information about choice grows gradually throughout the trial whereas information about stimulus increases stepwise early during the trial (Fig 4G, middle).

Finally, we asked whether this information was “local” (contained in individual whisk cycles; Isett et al., 2018) or continuous (integrated over the trial). We removed information from “sampling whisks” (those large enough to reach the shapes at their closest position) by setting the spike count to zero on those whisks, again using trial balancing to disentangle stimulus and choice. This largely abolished the encoding of stimulus but preserved the encoding of choice (Fig 4G, right), demonstrating that barrel cortex only transiently carries stimulus information during sampling whisks but encodes choice on longer timescales.

In sum, barrel cortex neurons respond to movements and contacts on fine timescales, and cognitive variables like choice can be decoded from these neurons on broad timescales. We next sought to understand the relationship of neural activity to choices.

### Distributed coding of sensorimotor and cognitive variables

To understand how individual neurons participate in the transformation of sensorimotor signals to decisions, we used regression to assess how they encoded local sensorimotor features and task-related features like choice. Specifically, we regressed the firing of each individual neuron onto each variable on individual whisk cycles using generalized linear models (GLMs; Fig 5A; Supplemental Fig 6A), similar to receptive field mapping by reverse correlation with natural stimuli (Park et al., 2014; Sharpee, 2013). Because neurons also encode movement and choice, this approach is needed to disentangle sensorimotor variables rather than simply attributing all responses to the presence of the shape (Fassihi et al., 2020).

**Figure 5.**
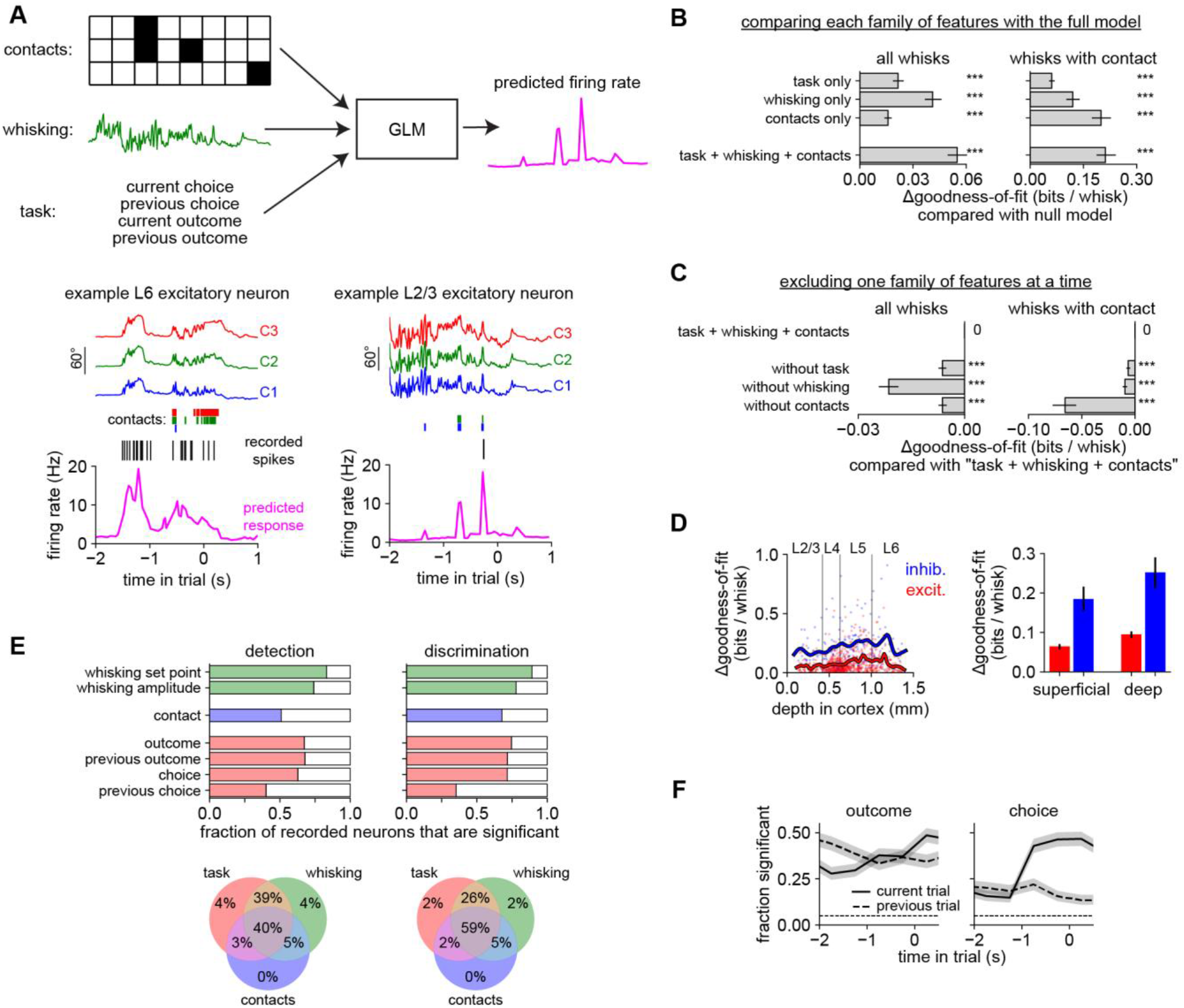
Heterogeneous coding of sensorimotor and task-related features. **A)** We used features from contacts (whisker identity), whisking (amplitude and set point), and task (choice and reward history) to train a GLM to predict neural responses on individual whisk cycles in held-out data. *Bottom left:* Example model predictions of firing rate (pink) for an example neuron (black raster: recorded spikes) given the position of each whisker (colored traces) and contacts (colored bars). This L6 neuron mainly responded to whisking, regardless of contacts. *Bottom right*: This L2/3 neuron mainly responded to contacts regardless of whisking. On this trial, it spiked only once. **B)** The goodness-of-fit (*i.e.*, ability to predict neural responses) of the GLM using features from the task, whisking, or contact families. Each feature family significantly improves the log-likelihood over a null model that used only information about baseline firing rate (p < 0.001, Wilcoxon test). The full model (“task + whisking + contacts”) outperforms any individual feature family. Similar results are obtained when testing on the entire dataset (*left*) or only on whisks with contact (*right*). n = 301 neurons during shape detection and n = 675 neurons during shape discrimination, pooled because the results were similar. **C)** The effect on goodness-of-fit of leaving out one family at a time from the full “task + whisking + contacts” model. Removing any of these feature families significantly decreases the goodness-of-fit, showing that each family contains unique information that cannot be obtained from the other families. **D)** Goodness-of-fit versus cortical depth (left) and grouped by cell type (right) in the “task + whisking + contacts” model. **E)** *Top:* Proportion of neurons that significantly encoded each variable during each task (p < 0.05, permutation test). *Bottom:* Venn diagram showing percentage of neurons significantly encoding features from task (red), whisking (green), and contact (blue) families during each task. In both tasks, a plurality of neurons encoded all three. <1% of neurons did not significantly encode any of the features. **F)** Proportion of neurons significantly modulated by the outcome or choice of the previous (dashed) or current (solid) trial during shape discrimination. Timescale defined in Fig 1C. Error bars: 95% confidence intervals, obtained by bootstrapping (B-D) or from a fit to a binomial distribution (F).

We treated the whisk as the fundamental unit of analysis rather than using arbitrary time bins because this granularity was useful for identifying behavioral strategies (Fig 3) and because contacts (Fig 2C) and spikes (Fig 4E) are highly packetized by the whisk cycle. Therefore we predicted total spike count on each whisk cycle for each neuron.

We again used model selection to quantify the importance of each feature for predicting neural responses. Different GLMs were trained on individual families of features—contact (“whisks with contact” as above), whisking (amplitude and set point), and task-related (choice and outcome of current and previous trial)— and their goodness-of-fit (*i.e.*, accuracy with which they predicted neural responses on held-out data) compared. Each family of features alone had explanatory power, and a combined “task + whisking + contacts” model surpassed any individual family (Fig 5B).

By individually dropping each family from the “task + whisking + contacts” combined model, we were able to assess whether explanatory power was unique to each family or instead shared across families due to their correlation. In each case, this significantly lowered the goodness-of-fit (Fig 5C), indicating that each family contained some unique information. Goodness-of-fit varied widely across the population but was in general highest in inhibitory and deep-layer neurons (Fig 5D).

Individual neurons could have been selective for specific features or broadly tuned for all features. We found that a plurality of neurons were significantly modulated by all three families (task, whisking, or contacts; Fig 5E). The results were strikingly similar across both discrimination and detection tasks. Thus, across behaviors, individual neurons in barrel cortex are broadly tuned for a mix of sensorimotor and task-related variables.

Finally, we investigated the dynamics of task-related variables over the course of the trial. Early in the trial, neurons encoded the outcome (rewarded or unrewarded) of the previous trial (Fig 5F), even though the previous trial could have been up to 12 s prior (*e.g.*, after an error timeout). This is surprising because the behavioral analysis revealed no effect of trial history on the mouse’s choice. Later in the trial, many neurons also encoded the choice on the current trial. This timecourse was similar to that of neural decoding (Fig 4G), which could not distinguish between coding for choice *per se* versus coding for the sensorimotor signals (contacts or whisking) that might correlate with choice. The GLM analysis disentangles these variables and demonstrates that, in addition to coding for sensorimotor variables, barrel cortex neurons also code for choice and outcome, persistently and on long timescales.

### Cell type-specific encoding of movement and contact

The tuning of individual neurons varied with cell type (excitatory or inhibitory) and laminar location (superficial or deep). The most prominent effect was that whisking strongly drove deep-layer inhibitory neurons (Fig 6A-C). Indeed, almost all (94 / 107 = 87.9%) inhibitory neurons in the deep layers were significantly excited by whisking (mean increase in firing rate: 23.9% per 10° of whisking amplitude). Excitatory neurons and superficial inhibitory neurons also encoded whisking, but were as likely to be suppressed as excited.

**Figure 6.**
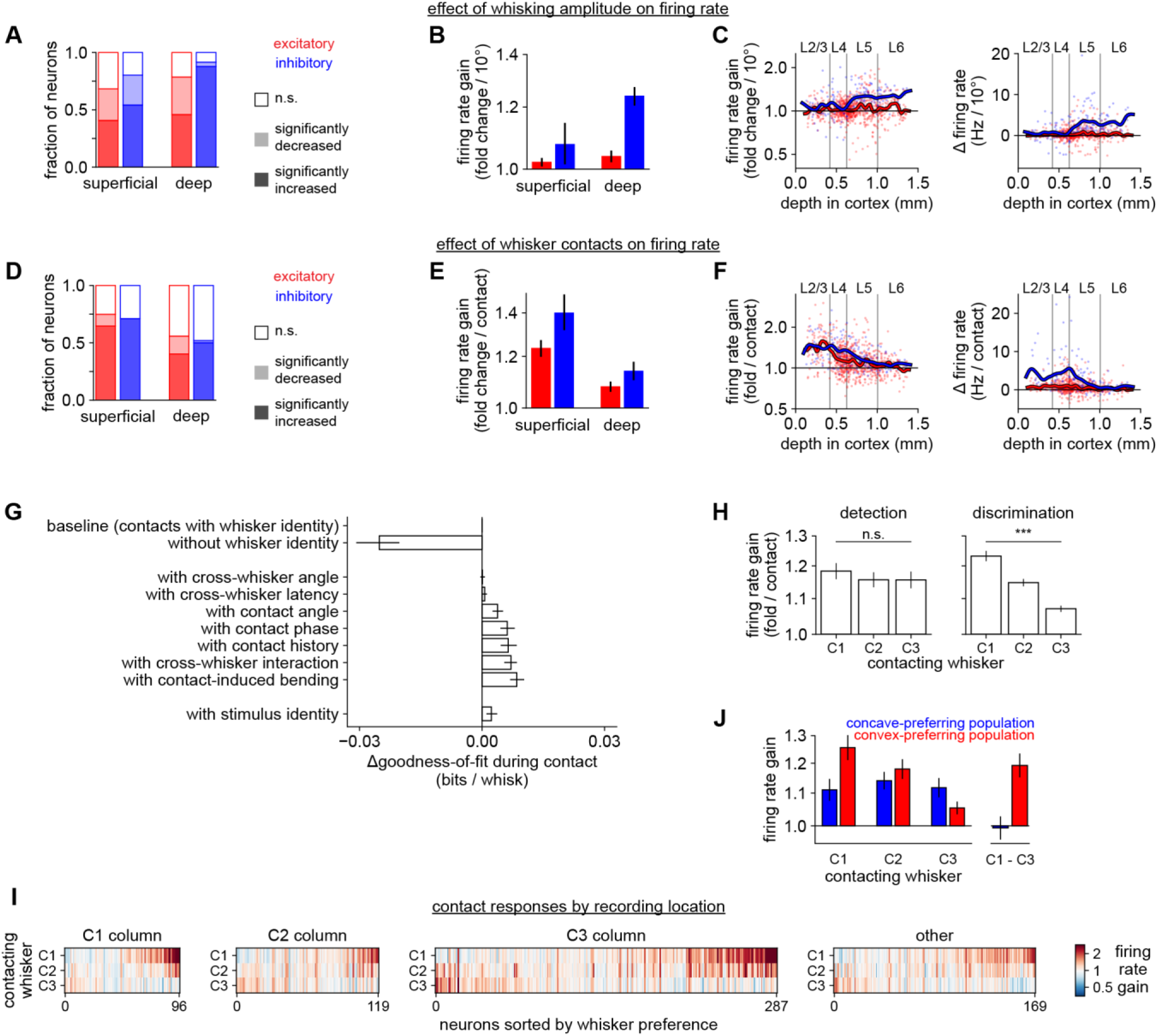
The formatting of contact responses facilitates decoding of shape identity. **A)** Proportion of neurons that are significantly excited (dark bars), suppressed (light bars), or unmodulated (open) by whisking amplitude, separately by cell type (p < 0.05, permutation test). In (A-C), n = 301 neurons during shape detection and n = 675 neurons during shape discrimination, pooled because the results were similar. **B)** Firing rate gain per each additional 10 degrees of whisking amplitude, grouped by cell type. Gain of 1.0 indicates no effect. **C)** Data in (B) for individual neurons plotted versus cortical depth. Blue and red lines: smoothed with a Gaussian kernel. *Left*: firing rate gain; *right*: change in absolute firing rate. **D-F)**Like panels (A-C), but for whisker contacts (averaged across C1, C2, and C3 whiskers). In (D-H), n = 235 neurons during shape detection and n = 675 neurons during shape discrimination. We excluded neurons for which too few whisker contacts occurred to estimate a response. **G)** Goodness-of-fit of models using different kinematic features about contacts, compared to the “task + whisking + contacts” model (top row, “baseline”). Removing whisker identity (second row) markedly decreases the quality of the fit. Adding additional kinematic parameters provides only a small increase in fit quality. The last row includes stimulus identity. **H)** Response to contacts of each whisker. *Left:* During shape detection, the population responds nearly equally to each whisker (p > 0.05; one-way ANOVA). *Right:* During shape discrimination, the population strongly prefers C1 contacts (p < 0.001). **I)** Responses to contacts made by each whisker for every recorded neuron during shape discrimination (n = 675), split by columnar location of the neuron. Tuning is diverse but neurons preferring C1 contacts are more common. **J)** The data in (H) broken into subpopulations that responded more during convex choices (n = 110; red) or concave choices (n = 76; blue), as assessed by the decoder analysis in Fig 4G. Neurons that fire more for convex choices respond much more strongly to C1 contacts than to C3 contacts, similar to the weights used by the optimized behavioral decoder (*cf.* Fig 3G). Logarithmic y-axis in B-C, E-F, H, and J. Error bars: 95% bootstrapped confidence intervals in B, E, and G; SEM in H and J.

In contrast, whisker contacts on the shapes more strongly excited superficial neurons, including both L2/3 and L4, relative to their baselines than those in deep layers (Fig 6D-F). Suppression by contact was less frequent than excitation in all cell types. Thus, movement and contact appear to have their greatest impact on the deeper and superficial layers, respectively.

### Contact responses are dominated by whisker identity, not finer sensorimotor parameters

We next asked which features of these contacts drove neurons and how this related to shape discrimination. Barrel cortex is arranged topographically with neurons in each cortical column typically responding to the corresponding whisker. Therefore preference for specific whiskers (somatotopy) was a clear candidate, but barrel cortex neurons are also tuned for multiple whiskers, contact force, whisking phase, cross-whisker timing, and global coherence, among other features (Brumberg et al., 1996; Drew and Feldman, 2007; Ego-Stengel et al., 2005).

Feature importance was assessed by comparing the goodness-of-fit of GLMs that had access to each feature. Whisker identity (which whisker made contact) was the most critical element determining neural firing (Fig 6G). The exact kinematics of contacts were less important. Of all the kinematic parameters we considered, contact force explained the most neural activity, but even this effect was relatively small compared to whisker identity.

We considered the possibility that some alternative kinematic feature that was not measured (*e.g.*, due to limitations in viewing angle or frame rate) might be driving neural activity. We therefore fit a model that also included the identity of the shape (concave or convex) on which each contact was made. If any unmeasured kinematic feature drove neural activity differently depending on the stimulus, this feature should capture some neural variability. Instead, it only slightly improved the model (Fig 6G), even less than including whisker bending (which did not strongly differ between the stimuli). This rules out, at least in a GLM framework, a latent variable that differentiates the stimuli and strongly drives neural activity.

Thus, during shape discrimination, contact responses are mainly driven by the identity of the whisker making contact. This was also the key feature for decoding stimulus and choice from the behavioral data, suggesting that contact responses might be dynamically modulated by task demands.

### Task-specific representation of contacts

Given that the identity of the contacting whisker was so critical for explaining neural responses, we examined each neuron’s tuning using the weights that the GLM assigned to each whisker. Neurons were spatially tuned, exhibiting whisker preferences (Supplemental Fig 6B,C). We did not select for responsive neurons, but rather included all neurons for which we had at least 10 contacts from each whisker.

During shape detection, the population of recorded neurons as a whole responded nearly equally to contacts made by C1, C2, and C3 (Fig 6H, left). Individual neurons could prefer any of the three whiskers, and in keeping with the somatotopy of barrel cortex, superficial neurons tended to prefer the whisker corresponding to their cortical column (Supplemental Fig 6D).

In marked contrast, we observed a widespread and powerful bias during shape discrimination: at the population level, neurons responded much more strongly to C1 contacts than to contacts by C2 or especially C3 (Fig 6H, right). Neurons preferring C1 were more prevalent in all cell types and in all recording locations, including the C2 and C3 cortical columns (Supplemental Fig 6E,F; individual neurons in Fig 6I). In an apparent violation of somatotopic organization, neuronal preference across columns was dominated by C1 regardless of the anatomical location. Because this preference was specific to the discrimination task, it could not be a trivial artifact of the shape stimuli or our analysis. Thus, whisker tuning was task-specific and strong enough to override somatotopy.

### Whisker-specific tuning explains the population choice signal

This somatotopic remapping during shape discrimination corresponds with the different weights assigned to each whisker by the behavioral classifiers (Fig 3G; C1 indicates convex). This correspondence suggests that neurons are retuned to C1 contacts in order to promote convex choices. It would have been equally plausible for neurons to prefer C3 contacts in order to promote concave choices, but this was not observed. This mirrors our behavioral observation (Fig 1G) that mice seemed to rely on a “convexity detection” strategy (*cf.* the “yes/no” tasks in Jang et al., 2009; Schulman and Mitchell, 1966).

We asked whether neurons’ choice preferences could be explained by their whisker tuning. Specifically, we assessed the tuning of two subpopulations of neurons preferring either concave or convex choices (*i.e.*, those assigned positive or negative weights by the decoder in Fig 4G). Indeed, the convex-preferring subpopulation strongly preferred C1 contacts (Fig 6J, red bars).

In summary, our neural encoder model (Fig 5-6) explains how the neural decoder (Fig 4G) was able to predict stimulus and choice: neurons were tuned for sensory input that the mouse had learned to associate with convex shapes. These representations were task-specific (Fig 6H) and could not be explained solely by simple geometrical aspects of the stimuli or whiskers. Indeed, the representations match weights used by the behavioral decoders to identify shapes. Our results link the tuning of individual neurons for fine-scale sensorimotor events to the more global and persistent representations of shape and choice. This bridging of local features to global identity is the essential computation of shape recognition.

## Discussion

In this study, we developed a novel head-fixed shape discrimination behavioral paradigm. Mice accomplished this task by comparing contacts made across whiskers. Barrel cortex neurons exhibited distributed coding of sensory, motor, and task-related signals. Deep inhibitory neurons encoded motion signals, and all neurons coded for contacts with a bias toward the whisker (C1) that preferentially contacted convex shapes. In mice performing shape detection, we observed similar coding of exploratory motion signals and of choice and outcome-related signals, but not the somatotopic bias. Thus, the barrel cortex relies partly on computational principles that are shared across tasks, and partly on task-specific coding that permits readout of behaviorally relevant variables.

### Behavioral decoding reveals sensorimotor strategies

Understanding neural mechanisms and computations begins with defining the subject’s behavioral goals and elucidating its strategy for achieving them (Krakauer et al., 2017; Marr and Poggio, 1976). Our approach was to measure many sensorimotor parameters about how mice interacted with the shapes and then to use behavioral decoding to predict the stimulus and choice from these data. This agnostic approach allowed us to rule out variables that contained little or no relevant information, identify from weights how animals interpreted informative variables, and form hypotheses for interpreting the neural data. We probed only one aspect of shape discrimination here, but this approach may easily be extended to other objects and tasks.

In two-alternative and go/nogo tasks stimulus and choice are correlated, especially when the subject’s accuracy is high. We disentangled stimulus and choice through trial balancing—overweighting incorrect trials so that in aggregate they are weighted the same as correct trials. This was crucial for interpreting the behavior. Some variables, such as contact count, were important for both stimulus and choice, indicating they were useful and actually used by the mouse. Others, such as contact angle, were more useful for predicting stimulus than choice, suggesting that mice did not (or could not) effectively exploit it. This effect is likely due to the incomplete information mice have about the instantaneous location of the whisker tips (Fee et al., 1997; Hill et al., 2011; Moore et al., 2015; Severson et al., 2019).

There are other approaches to disentangling stimulus and choice: separately fitting correct and incorrect trials, comparing stimulus prediction with choice, *etc.* (Campagner et al., 2019; Isett et al., 2018; Waiblinger et al., 2018; Zuo and Diamond, 2019a). We prefer trial balancing because it jointly optimizes over correct and incorrect trials. Regardless of the method chosen, the correlation between stimulus and choice must be considered, both in behavioral strategy analysis and in neural decoding.

### Mice compare contact counts across whiskers to discriminate shape

Shape discrimination fundamentally differs from pole localization and texture discrimination because it explicitly requires integration over different regions of space. Thus, comparing input across whiskers was a reasonable strategy for mice to pursue. Although mice can perform other tasks better with multiple whiskers (Carvell and Simons, 1995; Celikel and Sakmann, 2007; Knutsen et al., 2006; O’Connor et al., 2010a), those cases likely reflect statistical pooling of similar information from multiple sensors as in our shape detection control task (Krupa et al., 2001).

Our results go beyond statistical pooling. We are not aware of any published examples of mice assigning opposite behavioral meaning to input from different nearby whiskers. This strategy more closely captures the way primates compare across fingers when grasping objects (Davidson, 1972; Thakur et al., 2008) and suggests key roles for cross-columnar integration in somatosensory cortex (Petreanu et al., 2012; Thakur et al., 2012).

For shape discrimination, the identity of the contacting whiskers was the most important feature determining both behavioral choice and neural responses. Although cross-whisker timing has been hypothesized to be an important parameter for shape discrimination (Benison et al., 2006; *cf.* primate fingertips in Johansson and Flanagan, 2009), it was uninformative about shape in our task. This may be because whisker mechanics (“floppiness”) during movement add too much variability to cross-whisker contact latency. It has also been proposed that the pattern of forces over the whisker array as they “grasp” an object could be informative about shape (Bush et al., 2016; Hobbs et al., 2016a). We did not observe whisker bending to be particularly informative, but finer measurements of the moments and forces at the follicle may shed light on this (Yang and Hartmann, 2016).

In sum, we find that whisker identity during contact alone to be the critical parameter for shape discrimination. We cannot exclude the possibility that mice employed this strategy because we had trimmed off the other rows of whiskers, but a similar observation has been made in freely moving rats with all of their whiskers (Hobbs et al., 2016b), albeit not during goal-directed behavior.

### Efficient motor exploration strategies simplify the sensory readout

Reflecting this difference in strategy, mice interacted with the shapes in a fundamentally different way than in many previous tasks. In our task, mice lightly tapped the stimuli with the tips of multiple whiskers simultaneously. This “minimal impingement” approach (Mitchinson et al., 2007) is likely the natural mode of the whisker system (Grant et al., 2009; Ritt et al., 2008). Multiple light touches could also engage adaptation circuits within the somatosensory pathway, enhancing their ability to perform fine discrimination (Wang et al., 2010). In contrast, mice locate and detect poles by contacting them with high enough force to cause substantial whisker bending (Hong et al., 2018; Pammer et al., 2013). This likely amplifies the corresponding neural signal, an efficient strategy for detection (Campagner et al., 2016; O’Connor et al., 2010b; Ranganathan et al., 2018) though perhaps more useful for nearby poles than for surfaces.

A common thread running through the entire literature of whisking behavior is that animals learn a motor exploration strategy optimized for the task at hand: targeting whisking to a narrow region of space to locate objects (Cheung et al., 2019; O’Connor et al., 2010a), rubbing whiskers along surfaces to generate the high-acceleration events that correlate with texture (Isett et al., 2018; Jadhav et al., 2009; Schwarz, 2016), or targeting contacts to specific whiskers in the present work. Thus, animals pursue a motor strategy that simplifies the sensory readout, *e.g.* to a threshold on spike count (O’Connor et al., 2013). Performance is consequently limited by errors in motor targeting rather than in sensory detection (Cheung et al., 2019). Similarly, humans learn efficient motor strategies for directing gaze and grasp (Gamzu and Ahissar, 2001; Yang et al., 2016b).

The challenge of these tasks may lie in learning a skilled action that enhances active perception rather than in drawing fine category boundaries through sensory representations as in classical perceptual learning. Behavior may thus be considered a motor-sensory-motor sequence combining purposive exploration and sensory processing to guide further actions (Ahissar and Assa, 2016).

### Timescales of integration in barrel cortex and beyond

At the sensory periphery neurons encode moment-to-moment sensory input whereas high-order brain structures encode object identity persistently and often with spatial invariance. Barrel cortex, an intermediate stage in this process, can be driven by extreme <5 ms synchrony (Bruno and Sakmann, 2006), the previous ~50 ms of sensory input (Bale and Maravall, 2018; Fassihi et al., 2020; Ramirez et al., 2014), whisker motor movements at ~100 ms (Fee et al., 1997), and even task-related features like reward over >1 s timescales (Lacefield et al., 2019).

Barrel cortex is thus well-situated to bind precise sensory information to longer-lasting internal states. Rather than generating decisions *per se*, its role may be to format sensorimotor information in a task-specific way, furnishing the results to downstream areas like secondary somatosensory cortex, primary motor cortex, parietal cortex, or striatum (Chen et al., 2013; Mohan et al., 2019; Sippy et al., 2015; Yang et al., 2016a; Zuo and Diamond, 2019b). In support of this formatting hypothesis, we find that neurons encode short-timescale events like contacts and whisks, but that decoders can also read out stimulus identity and choice over the course of the trial.

Previous trial outcome, which had little effect on the mouse’s choice, prominently modulated neural activity. This could reflect arousal or a long-lasting effect of reward (Lacefield et al., 2019; McGinley et al., 2015; Vinck et al., 2015). Persistent representations of choice and outcome have been observed in prefrontal, parietal, and motor cortex; the thalamus; and indeed throughout the brain (Akrami et al., 2018; Hattori et al., 2019; Lavzin et al., 2020; Nogueira et al., 2017; Tsutsui et al., 2016; Waiblinger et al., 2018). Recent work in artificial neuronal networks suggests that distributed outcome signals may be a general principle of reinforcement learning (Dabney et al., 2020).

### Motor signals in barrel cortex

In natural behavior, active sensing is the norm: animals explore by moving their heads, eyes, and ears and by sniffing, chewing, or grasping objects. Motor signals should perhaps be expected in sensory areas because they provide context for interpreting sensory input. We found that movement was generally the largest signal in barrel cortex activity, stronger even than responses to whisker contacts. Recent studies have variously found that barrel cortex neurons respond to whisking onset (Muñoz et al., 2017; Yu et al., 2016), that whisking phase modulates contact responses (Curtis and Kleinfeld, 2009; Hires et al., 2015), or that whisking simply has mixed effects on neuronal firing (Ayaz et al., 2019; O’Connor et al., 2010b; Peron et al., 2015). Technical limitations of whisker tracking perhaps explain these disparate results (Krupa et al., 2004).

In resolving this, we found that multiple features of whisking are encoded simultaneously: whisking set point and amplitude (global signals shared across whiskers) and whisking pose (the relative angle of each whisker to the others, *cf.* hand posture in Goodman et al., 2019). This encoding was widespread but with a strong cell type-specific bias: inhibitory neurons in the deep layers were robustly and consistently excited in proportion to whisking amplitude. This is consistent with studies of parvalbumin-positive neurons in layer 4 and 5B/6 (Yu et al., 2016), as well as in other classes of inhibitory interneurons throughout all layers (Muñoz et al., 2017; Yu et al., 2019). However, these studies focused on responses around whisking onset rather than the continuous and graded encoding of whisking that we describe here.

One theory is that whisker motion should be encoded in inhibitory signals so that the brain can predict and account for the sensory consequences of movement (Yu et al., 2016), as in the auditory cortex (Schneider et al., 2018). The inhibitory neurons of the deep layers receive direct input from primary motor cortex (Kinnischtzke et al., 2014) and can potently suppress the entire cortical column (Bortone et al., 2014; Frandolig et al., 2019). A subtraction could account for simple stimuli, but increasingly complex stimuli like shapes may require mixed selectivity and distributed coding of sensorimotor signals (Rigotti et al., 2013). This perhaps explains the diverse tuning for whisking that we observed in excitatory neurons.

More generally, whisker motion signals may be analogous to the preparatory saccade signals identified in visual cortex. The brain may exploit the synchronization and discretization of sensory signals to make judgments about the external world. Like whisking, saccades are a motor action directed toward collecting information, and the cortex predicts the resulting change in sensory input (Steinmetz and Moore, 2010).

### Barrel cortex formats contact representations for reading out shape

At first glance, the whisker system may appear to be a labeled line system due to its somatotopic organization at the level of the brainstem, thalamus, and cortex. Indeed, neurons in thalamorecipient layer 4 typically respond best to stimulation of an anatomically corresponding whisker. However, outside of L4 the preference for any particular whisker is much weaker (Brecht et al., 2003; Clancy et al., 2015; Jacob et al., 2008; De Kock et al., 2007; Peron et al., 2015; Pluta et al., 2017; Ramirez et al., 2014), and indeed attending to whisker input decreases somatotopy (Wang et al., 2019).

Rather than maintaining a labeled-line code, the barrel cortex may encode multi-whisker sequences, a map of scanned space, or entire tactile scenes (Laboy-Juárez et al., 2019; Pluta et al., 2017; Vilarchao et al., 2018; reviewed in Estebanez et al., 2018). Similarly, auditory cortex is now thought to encode high-level sound features rather than strict tonotopy (Bandyopadhyay et al., 2010; Carcea et al., 2017; Rothschild et al., 2010). Ethologically, integrating information across sensors would seem far more useful than maintaining in higher-level areas a strict segregation based on peripheral organization. In keeping with this, we observed a substantial diversity across individual neurons in their tuning for contacts by each whisker, and an over-representation of behaviorally relevant whiskers during shape discrimination.

### Task-specific coding for efficient sensorimotor identification

We suggest that the barrel cortex learns to code preferentially for the sensory features that are most relevant for the animal’s goals (Ramalingam et al., 2013). In shape discrimination, mice learned to compare the space sampled by the C1 whisker with the space sampled by the C3 whisker. One way to “detect convex” shapes is to preferentially enhance C1 contacts (overrepresented on convex shapes) and suppress C3 contacts, essentially implementing a cross-whisker subtraction. Thus the neural responses to contacts are reweighted to permit the detection of convexity. This computation relies on both a motor strategy (targeting contacts on each shape to specific whiskers) and a neural coding mechanism (enhancing responses to contacts on specific whiskers). Further studies are needed to assess the degree to which neural mechanisms depend on local plasticity versus descending input from higher areas.

One intriguing possibility is that these computations may be related to efficient coding. Within the timescale of a single trial, mice may refine increasingly accurate models of the shape, as they learn to predict future sensory input from earlier input in the trial. In turn, they could adopt exploratory motion strategies to test their current prediction. These prediction signals are thought to be represented by different cortical layers (reviewed in Adesnik and Naka, 2018), and deficits in predictive coding have been hypothesized to explain the differences in the autistic and schizophrenic brains (Keller and Mrsic-Flogel, 2018; Robertson and Baron-Cohen, 2017).

The superficial and deep layers of cortex can encode sensory stimuli independently (Constantinople and Bruno, 2013) but they can also strongly interact (Pluta et al., 2019). We observed stronger touch responses in the superficial layers and stronger whisking responses in the deep layers, potentially useful for simulating the effects of motor exploration (Brecht, 2017). The superficial layers may also encode mismatch signals like unexpected contacts (Keller et al., 2012; although *cf.* Ayaz et al., 2019). Further work will be necessary to determine how and when this translaminar circuit implements active sensation.

Although the details of these effects are specific to this task and stimulus geometry, we suggest that analogous computations in other brain areas and species could implement object recognition by comparing input across different sensors in the context of exploratory motion. Recent results have demonstrated an unexpectedly widespread coding for motion across the brain (Musall et al., 2019; Stringer et al., 2019). These motion signals could be critical for interpreting sensory input in the context of behavioral state. The common structure of cortex across regions of disparate functionality (Douglas and Martin, 2004) may be a signature of this common computational goal.

## Author Contributions

CR, SF, and RB conceived of the experiments and analytic approach. CR developed the behavior with help from EG and BCP who also assisted in training the mice. CR and EG performed videography. CR built the electrophysiological recording rig, performed all electrophysiological recordings, and coded the videographic and electrophysiological analyses. RN and SF provided insight and guidance on the theoretical concepts and analytic approach. CR and RN analyzed data and CR generated the final versions of analyses and figures presented here. CR, RN, SF, and RB together decided how to interpret the data. CR wrote and RN, SF, and RB edited the manuscript.

The authors declare no competing interests.

## Acknowledgements

We thank Akash Khanna and Philip Calafati for help with mouse training; Drew Baughman for histological assistance and lab management; Clay Lacefield for early assistance with behavioral apparatus; Jung M Park for assistance with building behavioral apparatus, mouse training, and perfusions; Richard Warren for help with whisker tracking algorithms; Caroline Bollen for assistance with microscopy; Brian Isett for advice on longitudinal acute recording technique; Michael Shadlen and Anita Devineni for comments on the manuscript; and the members of the Center for Theoretical Neuroscience for insightful suggestions.

Support was provided by NINDS/NIH (R01NS094659, F32NS096819, and U01NS099726); NeuroNex (DBI-1707398); the Gatsby Charitable Foundation (GAT3419); a Kavli Institute for Brain Science postdoctoral fellowship (to CR); and a Brain & Behavior Research Foundation Young Investigator Award (to CR).

The content is solely the responsibility of the authors and does not necessarily represent the official views of the NIH.

## STAR Methods

### Resource Availability

#### Materials Availability

This study did not generate any unique reagents.

#### Data and Code Availability

Upon publication of this manuscript, all data and code necessary to generate the results presented here will become publicly available at github.com and crcns.org.

### Experimental Model and Subject Details

We report data here from 17 adult mice (11 females and 6 males) of the C57BL6/J strain bred in the Columbia University animal facilities. 11 mice were used for shape discrimination, 5 for shape detection, and an image was used from 1 mouse from a different, anatomical study (Fig 4A). Of these, one shape discrimination mouse was discarded from all video analysis because of poor video quality, but was still used for behavioral performance data in Supplemental Fig 1A.

Mice in our colony are continuously backcrossed to C57BL6/J wild-type mice from Jackson Laboratories. Some mice expressed Cre, CreER, Halorhodopsin, Channelrhodopsin2, and/or EGFP for ongoing and unpublished studies. We noted no difference in the results regardless of the genes expressed and therefore pooled the data here. Mice were group-housed (unless they did not tolerate this) and lived in a pathogen-free barrier facility. All experiments were conducted under the supervision and approval of the Columbia University Institutional Animal Care and Use Committee.

### Method Details

#### Surgeries

Mice were implanted with a custom-designed stainless steel headplate (manufactured by Wilke Enginuity) between postnatal day 90 and 180. They received carprofen and buprenorphine and were anesthetized with isoflurane throughout the stereotaxic procedure. Using aseptic technique, we removed the scalp and fascia covering the dorsal surface of the skull. We then positioned the headplate over the skull and affixed it with Metabond (Parkell).

After behavioral training (see below), some mice underwent another procedure to permit electrophysiological recording. First, we used a dental drill to thin the cement and skull over barrel cortex, rendering it optically transparent, and coated it with cyanoacrylate glue (Vetbond). We used intrinsic optical signal imaging (described below) to locate the cortical columns of the barrel field corresponding to the whiskers on the face. We then used a scalpel (Fine Science) to cut a small craniotomy directly over the columns of interest. Between recording sessions, the craniotomy was sealed with silicone gel (Dow DOWSIL 3-4680, Ellsworth Adhesives) and/or silicone sealant (Kwik-Cast, World Precision Instruments).

#### Intrinsic signal optical imaging

Individual barrel-related cortical columns were located with intrinsic imaging. While the mice were anesthetized with isoflurane, individual whiskers were deflected one at a time by a piezoelectric stimulator (8 pulses in the rostral direction at 5 Hz, with ~30 s between trains). We used custom software written in LabView (National Instruments) to acquire images of the cortical surface through the transparent thinned skull under a red light source with a Rolera CCD camera (QImaging). Videos were averaged over 20-60 trains of pulses. We repeated this procedure for the C1, C2, and C3 whiskers to locate the region of maximal initial reflectance change corresponding to each.

#### Behavioral apparatus

The behavioral apparatus was contained within a black box (Foremost) with a light-blocking door. It was built with posts (Thorlabs) and custom-designed laser-cut plastic pieces on an aluminum bread board (Edmund Optics, Thorlabs, or Newport). A stepper motor (Pololu 1204) rotated a custom-designed curved shape 3D-printed with ABS plastic (Shapeways) into position, and a linear actuator (Actuonix L12-30-50-6-R) moved it within reach of the mouse’s whiskers. Rewards (~5 μL of water, chosen based on the mouse’s weight and how many trials it typically completed) were delivered by opening a solenoid valve (The Lee Co. LFAA1209512H) that allowed water to flow to the mouse from a reservoir to a thin stainless steel tube (McMaster).

An Arduino Uno, in communication with a desktop computer over a USB cable, controlled the motors. It also monitored licking by sampling beam breaks of the mouse’s tongue through infrared proximity detectors (QRD1114, Sparkfun) or capacitive touch sensors (MPR121, Sparkfun) in front of and slightly to the left or right of the mouse’s mouth, inspired by a published two-choice design (Guo et al., 2014). Between trials only, the Arduino activated a white “house light” (LE LED; Amazon B00YMNS4YA) that prevented mice from fully dark-adapting, preventing the use of visual cues. A computer fan (Cooler Master; Amazon B005C31GIA) continuously blew air slowly over the shape such that the mouse’s nose was upwind from the shape, preventing the use of olfactory cues. We never observed mice exploiting auditory or vibrational cues from the motors and thus no masking noises were necessary.

At a fine timescale the trial structure was controlled by the Arduino using a custom-written sketch. At the level of individual trials, the desktop PC chose the stimulus and correct response and logged all events read from the Arduino to disk using custom Python code. The training parameters for each mouse were stored in a custom-written django database and updated manually or semi-manually by the experimenters depending on each mouse’s progress.

#### Behavioral training

Throughout, the mice were denied access to water in the home cage and learned to receive their water during behavioral training. We closely monitored their water intake, weight, and general health to ensure they did not become dehydrated. *Ad libitum* water was provided if necessary to ensure health.

Mice were trained to perform either the shape discrimination or detection tasks using a process of gradual behavioral shaping described below. Some mice were additionally trained to discriminate flatter, more difficult shapes.

1. “Lick training.” Mice initially learned to lick to receive water. They were advanced through each step of this stage only once they learned to receive sufficient daily water from the apparatus. First, they were placed in the apparatus without head-fixing and allowed to drink freely from the water pipes, which rewarded every lick. Next, we head-fixed the mice directly in front of a single lick pipe and rewarded every lick. Finally, mice were presented with two lick pipes (left and right) and learned to lick alternately from each of them, first in blocks of ten licks and gradually decreasing to a single lick on each side. This stage required 12.5 sessions on average.
2. “Forced alternation”. We introduced the complete trial structure for the first time, presenting shapes and rewarding the mouse only for correct responses and punishing it with a timeout for incorrect responses. During this stage the shape on each trial was not random; instead, mice were repeatedly presented with the same shape trial after trial until it gave the correct response. After a correct response, the other stimulus was presented. Thus, mice could perform at 100% by alternating responses from trial to trial. The timeout was initially 2 s and then increased to 5 s and finally 9 s as the mice became accustomed to it. This stage required 11.3 sessions on average.
3. “Stimulus randomization with bias correction”. During this stage, stimulus identity was randomized on each trial and only presented at the closest position. Each session began with 45 trials of “forced alternation” to ensure that mice were able to lick both directions. After that, trials were generally random. The software continuously monitored their performance for biases; when a strong bias was detected, it stopped presenting trials randomly and began presenting trials designed to counteract the bias. For instance, if mice responded on the left ≥20% more than on the right, the software would deliver only right trials. Alternatively, if the mice showed a significant perseverative bias (ANOVA “choice ~ stimulus + side + previous_choice”, p < 0.05 on previous_choice), the software would deliver “forced alternation” trials. Critically, we only ever analyzed truly random trials from the session. Non-random trials were used only for behavioral shaping and were discarded from behavioral and neural analyses.
4. “Range of positions”. We now presented shapes at the first 2 positions (close and medium) and then all 3 positions (close, medium, and far). Position was randomized across trials. The same automatic training and bias-prevention procedures as before were used.
5. “Flatter shapes”. Some mice were now presented with flatter shapes as well as the shapes of the original curvature. Other mice skipped this stage and were never presented with flatter shapes.
6. “Whisker trimming”. We gradually trimmed whiskers off the right side of the face: first we trimmed the A and E rows, then the B row, then the D row. After any trimming, we allowed mice to recover to high performance before trimming additional rows. We retrimmed previously trimmed whiskers as necessary to ensure they could not reach the shapes. Stages 3-6 required a total of 109.1 sessions on average.

Sometimes it was necessary to return mice to an earlier stage of training temporarily to facilitate learning (*e.g.*, reducing the number of positions at which the shapes were presented or returning to “forced alternation” trials only). Mice that successfully progressed through all stages of the training procedure— those who could identify both shapes at all three positions with only the C-row of whiskers—were deemed fully trained. We only took high-speed video or neural recordings from fully trained mice.

#### Videography

For videography and electrophysiology, we moved the behavioral setup to a different light-blocking box mounted on a vibration-isolating air table (TMC). We took video of fully trained mice using a high-speed camera (Photonfocus DR1-D1312IE-100-G2-8) with a 0.15 ms exposure time to prevent motion blur. We used a lens with a 25 cm focal length (Fujinon HF25HA-1B) to prevent “fisheye” distortion. An aperture (F-stop) of approximately 6.0 optimized depth of field.

We designed and built a custom infrared backlight with a 7×8 grid of high-power surface-mount infrared (850 nm) LEDs (Digikey VSMY2853G) soldered to a custom-designed PCB (manufactured by OSH Park) that allocated power to each LED through current-limiting resistors. Diffusion paper mounted above the LEDs homogenized the light. The backlight was placed below the mouse and pointed toward the camera so that the whiskers would show up as high-contrast black on a white background. The Arduino pulsed this backlight off for 100 ms at the beginning of each trial, allowing us to synchronize the behavioral and video data. We used Matlab’s Image Acquisition Toolbox to store the video data to an SSD.

#### Electrophysiology

To record neural activity, we head-fixed the mouse in the behavioral arena as usual and removed the temporary sealant over the craniotomy. We lowered an electrode array (Cambridge Neurotech H3) using a motorized micromanipulator (Scientifica PatchStar), noting its depth at initial contact and at final position. We used an OpenEphys acquisition system with two digital headstages (Intan C3314) to record 64 channels of neural data at 30 kHz at the widest possible bandwidth (1 Hz to 7.5 kHz). The backlight sync pulse was acquired with an analog input to synchronize the neural, behavioral, and video data.

We used KiloSort (Pachitariu et al., 2016) to detect spikes and to assign them to putative single units. Single units had to pass both subjective and objective quality checks. First, we used Phy (Rossant et al., 2016) to manually inspect every unit, merging units that appeared to be from the same origin based on their amplitude over time and their auto- and cross-correlations. Units that did not show a refractory period (*i.e.* a complete or partial dip in the auto-correlation within 3 ms) were deemed multi-unit and discarded. Second, single units had to pass all of the following objective criteria: ≤5% of the inter-spike intervals less than 3 ms; ≤1.5% change per minute in spike amplitude; ≤20% of the recording at <5% of the mean firing rate; ≤15% of the spike amplitude distribution below the detection threshold; ≤3% of the spike amplitudes below 10 μV; ≤5% of the spikes overlapping with common-mode artefacts.

We identified inhibitory neurons from their waveform half-width, *i.e.* the time between maximum negativity and return to baseline on the channel where this waveform had highest power. Neurons with a half-width below 0.3 ms were deemed narrow-spiking and putatively inhibitory. We measured the laminar location of each neuron (using the boundaries in Hooks et al., 2011) based on the manipulator depth and the channel on which the waveform had greatest RMS power. Neuron in L1 or the cortical subplate were discarded from this analysis because they were difficult to sort and showed variable properties across mice.

#### Histological reconstruction of recording locations

We used a camera mounted on a surgical microscope to take a picture of the area around barrel cortex on every session from the time of intrinsic signal imaging to the end of the experiment. We aligned all of these images with each other using the TrakEM plugin (Cardona et al., 2012) in Fiji using surface vasculature. These images, referenced to individual barrel column locations determined by intrinsic signal imaging, were used to guide the placement of the craniotomy and the electrode. We also photographed and aligned images of the location of the implanted electrode array each day.

On the last day, we inserted a glass pipette coated with DiI (Sigma-Aldrich 468495) into the barrel field twice to leave two landmarks, one anterior and one posterior, which were also photographed and aligned. At the conclusion of the experiment, we deeply anesthetized the mice with pentobarbital, transcardially perfused them with 4% paraformaldehyde, and removed the brain for histological processing.

The left hemisphere was sectioned tangentially to the barrel field using a Vibratome or freezing microtome to cut 50 or 100 μm sections. We stained for barrels with fluorescently conjugated streptavidin and imaged the sections on an epifluorescent microscope to reveal the location of the barrels and the DiI landmarks. In this way we confirmed the exact location of each recording site with respect to both the anatomical and functional barrel map.

### Quantification and statistical analysis

#### Statistics

Throughout this manuscript, “*” indicates p < 0.05; “**” indicates p < 0.01; “***” indicates p < 0.001; and “n.s.” indicates “not significant”.

To non-parametrically estimate the width of certain non-normal distributions, we used “bootstrapped confidence intervals”. This means resampling the data with replacement 1000 times, taking the average of each resampled dataset, and then taking the interval that spans the central 95% of this distribution of averages across resampled datasets.

#### Whisker video analysis

We used a lightly modified fork of the ‘pose-tensorflow’ package (Insafutdinov et al., 2016; Pishchulin et al., 2015) to train and use a deep convolutional neural network to identify and track whiskers in the video. This network is based on Resnet (He et al., 2015) and is the same “feature detector” network incorporated into the first version of DeepLabCut (Mathis et al., 2018). We generated an initial training set using the software ‘whisk’ (Clack et al., 2012) to track individual whiskers and custom semi-automated code to classify them.

Eight equally spaced points along each tracked whisker were provided as the “joints” for the neural network to identify. We iteratively improved the neural network by evaluating it on new frames, choosing difficult frames from the result, semi-automatically improving the labels, swapping in the results from ‘whisk’ as necessary, and then using this new training set to train a new version of the network. Whiskers of below-threshold confidence or below-threshold smoothness at any joint were discarded. We optimized these thresholds with a cross-validated grid search.

Sessions with inaccurate labeling were discarded: we required that every whisker be labeled in ≥95% of the frames, that ≤2% of the contact events contained even a single frame with a missing label, and that the arcs traced out over the entire session by the whisker contained no discontinuities or jumps suggestive of tracking errors. In the remaining well-traced sessions we interpolated whiskers over any missing frames.

We identified the shape stimulus in each frame by thresholding and segmenting the frame and selecting the segment of the appropriate size and location. We identified contacts on the shape based on proximity (≤10 pixels Cartesian distance) between the tip of each whisker and the edge of the shape.

To estimate each whisker’s bending moment, we first fit a spline through its 8 identified joints and used the “measure” function of ‘whisk’ to estimate curvature (κ). κ is the rate of change of direction of the whisker at each point along its length, *i.e.* the reciprocal of the radius of curvature at that point, and is measured in units of m^−1^. ‘whisk’ averages κ over the entire length of the traced whisker and we followed this convention. For comparison with other studies, we note that 1 m^−1^ is equal to 0.001 mm^−1^ due to this reciprocal. κ = 0 for a straight line. In our study, κ > 0 for a whisker pushing into a shape and κ < 0 for the reverse curvature, typically encountered while detaching from the shape.

To register all videos within a common reference frame for visualization (Fig 3I-K), we extracted the location of the shape edge at each location (close, medium, or far). Because we knew the exact distance between edges in reality, we used the vector between adjacent locations in the image to measure the angle and scale for that particular video. After compensating for this angle and scale, we used the peak in the 2D cross-correlation to find the offset that best aligned the videos with each other.

#### Decomposition of individual whisks

We defined the whisker’s angle as the Cartesian angle between base and tip. We decomposed the whisking signal into individual whisk cycles with the Hill transform (Hill et al., 2011). Briefly, we bandpassed the data from 8 to 50 Hz and applied the Hilbert transform to extract phase. Peaks and troughs were defined as frames where the phase crossed zero or pi. We defined set point as the angle of each whisker at the trough of each whisk cycle, and amplitude as the angular difference between peak and trough on each cycle for the C2 whisker. The whisking amplitude was very consistent across whiskers, so we used the amplitude of the C2 whiskers cycle only throughout. In contrast the relative set point of each whisker could vary, so we used the set point of each as regressors in the neural GLM analysis. To smooth these amplitude and set point parameters, we applied a triangular window that weighted one cycle before and after half as much as the current cycle.

To identify sampling whisks (those large enough to reach the shapes if they had been at their closest position), we aligned the frames to the response window and found the convex hull of the edges of the shape (i.e., the boundary of closest points to the whisker pad) versus time from the response window. A “whisk without contact” was one on which the whiskers crossed this boundary. This could happen if, for instance, the C3 whisker investigated the space where the medial portion of the closest concave shape would be, but actually a convex shape was present or a concave shape at a further position (example: Fig 3A). A “whisk with contact” is any whisk on which contact was made. Sampling whisks are defined as either “whisks with contact” or “whisks without contact”. All other whisks (non-sampling whisks) are those which did not cross the convex hull described above and did not make contact with the shapes. Not all trials contained contacts, but the vast majority of trials included at least one sampling whisk.

#### Lick rates (Fig 1D)

We recorded the times of all licks, even those before the response window that had no effect on the trial outcome. In rare cases our detector recorded a single lick as many licks (the “switch bouncing” effect) and so for analysis we binned licks in 100 ms bins and discarded any surplus licks above one per bin.

To plot the rate of correct or concordant licks, we calculated the rate of licking on each side on every trial and defined each lick as “correct/incorrect” depending on whether it matched the correct side, and as “concordant/discordant” depending on whether it matched the direction of the choice lick (*i.e.*, the first lick in the response window, which determined trial outcome). We then meaned the lick rates for each trial type (correct, incorrect, concordant, discordant) within each mouse. Finally we divided the rate of correct licks by the rate of all licks, and the rate of concordant licks by the rate of all licks, to generate the results plotted in Fig 1D.

#### Behavioral decoding analysis (Figure 3)

We first selected only trials in which the mouse responded within the first 0.5 s of the response window in order to ensure that behavior was roughly aligned across trials. This procedure excluded only a small fraction of trials. In some sessions we used optogenetic stimulation for separate studies; any trial with optogenetic stimulation was discarded from all analysis in this manuscript. In some sessions we also presented flatter shapes (performance data: Supplemental Fig 1A) but for behavioral decoding and all neural analyses we discarded any trials with the flatter shapes.

We then extracted all whisking and contact data from each trial from −2.0 to +0.5 s of the opening of the response window and obliviated (set to zero or the mean value) all data after the time of the choice lick to ensure that only pre-choice activity was included in the analysis. Each feature was measured on every individual whisk (e.g., presence of contact, cross-whisker latency within that contact, interaction terms for multiple-whisker contact; complete list in Supplemental Table 1). We then aggregated each feature within 250 ms bins locked to the response window opening, so that trials with different numbers of whisks could be directly compared. Most features were aggregated by meaning within the bin, but count-related features (like contact count) were aggregated by summing within the bin. Finally we concatenated some task-related features like previous choice and previous outcome that did not depend on the whisk cycle.

If a feature was not defined for a time bin (for instance, cross-whisker contact timing and contact-induced bending have no meaning if no contacts occurred), it was left as null (NaN). Because these parameters were only measured during contacts, they implicitly contained information that a contact had occurred. Specifically, they were null at all times other than during contact. During feature standardization (described below), we ensured that these features could have no effect on the coefficients or fit when they were null. The net result of this procedure is that these features could only be informative conditioned on the presence of a contact. This permits their interpretation as modulating the information gleaned by the mouse about each contact, above and beyond the mere presence of a contact *per se*.

The result of this feature selection process was 725 features per trial, some of which (e.g., contact count) depended on time bin and some of which (e.g., previous choice) did not. For each session, we standardized all features by scaling them to zero mean and unit standard deviation. At this point we imputed null (NaN) features with zero, so that they could not affect the coefficients obtained. We used the same procedures to fit individual features (Fig 2B) or combinations of features (Fig 2C).

##### Cross-validation scheme

Each session was fit separately. We grouped the trials into 4 separate “strata”, with one stratum for each combination of choice and stimulus (concave/convex for discrimination; something/nothing for detection). We split the data into 7 “folds” for cross-validation, equally sampling trials from each stratum. Each trial was in the “testing” set for one fold, the “tuning” set for one fold, and the “training” set for five folds. For each fold, we fit a logistic regression model (‘sklearn.linear_model.LogisticRegression’) on the training set over a range of different regularization parameters. We then evaluated the model on the held-out tuning set and chose the regularization that optimized classifier accuracy over all sessions. Finally we evaluated the model with the chosen regularization on the doubly held-out testing set and took that score as the model’s overall accuracy.

To analyze the weights of the classifier for the session as a whole, we averaged the weights across folds. To analyze the prediction on an individual trial, we used the classifier for which that trial was in the testing (doubly held-out) set. Because each trial was in the testing set in exactly one fold, there was only one unique prediction per trial.

##### Trial balancing to decorrelate stimulus and choice

We weighted every trial inversely to its prevalence in the dataset, using the ‘sample_weight’ argument. Essentially, if correct trials were three times more common than errors, then we weighted each individual error three times as much, so that the total weight assigned to correct and incorrect trials was equal. This balancing effectively decoupled the correlated target labels stimulus and choice. We validated this approach by comparing the results to other balancing schemes like undersampling to the size of the smallest stratum or over-sampling by bootstrapping (data not shown).

##### Aggregation

To aggregate results across mice (*e.g.,* Fig 3D,F) we averaged the accuracy of the classifier across sessions within each mouse first. The sample size for error bars and statistical tests was then equal to the number of mice.

To plot the weights of the classifier in Fig 3G, we first averaged the weights over time for simplicity. Because the coefficients plotted in Fig 3G are related to contact counts, we multiplied the coefficients by the standard deviation of the corresponding column in the feature matrix before standardization. This effectively reverse the normalization, and puts the coefficient in more-interpretable “per contact” units rather than “per standard deviation of contact count” units. This was for visualization only and did not affect the results.

To plot the evidence in Fig 3K, we applied the weights of the decoder to each individual whisk cycle and meaned this evidence over all whisks with a peak within that spatial bin. For visualization in this panel, we used a model that incorporated the peak angle of whisks without contact.

#### Neural decoding analysis (Fig 4G)

To decode stimulus and choice from neural activity, we used a resampling/bootstrapping approach to combine neural data across sessions and mice. First the trials were split into five equally sized “folds”, one of which was the “test fold” and the rest “train folds”. No tuning set was necessary because we fixed the regularization at 1.0 in this analysis. For each shape (concave or convex), we randomly chose a single trial with that shape from the test fold in each session. We concatenated all of the neural data from those trials into a “pseudopopulation” as if all the neurons had been recorded simultaneously. We then repeated this process 30 times to construct 30 pseudotrials of the test fold. Then, we repeated the process for the train folds, to generate 120 pseudotrials of the train folds. By construction, the same trial could never be included in both the test and train folds.

The classifier was trained on the train fold and evaluated on the test fold. Because correlations can have a strong impact on the amount of information encoded by a neuronal population (Nogueira et al., 2020), we maintained the correlation structure between simultaneously recorded neurons. Specifically, for each pseudotrial we sampled the same trial from each simultaneously recorded neuron. The entire process was repeated 100 times to generate the bootstrapped confidence intervals displayed in the plot.

We call the procedure above the “naive” approach because it does not balance hits and errors; hence, it confounds stimulus and choice. This naive approach is used in the left panel of Fig 4G. We also used a “balanced” approach to disentangle stimulus and choice in the middle and right panels of Fig 4G. Specifically, we first divided all the trials into 4 strata (concave hit, concave error, convex hit, convex error) instead of the 2 strata (concave or convex) used in the naive approach. We then repeated the same resampling approach to draw pseudotrials from each of the 4 strata. This ensures equal weighting of correct and incorrect trials; hence, it is balanced. We used disjoint train and test folds just as in the naive approach.

In all cases, to train the classifier we first standardized the firing rate of each neuron in the pseudopopulation to zero mean and unit variance. We provided these normalized firing rates to a classifier (‘sklearn.linear_model.LogisticRegression’) and trained it to predict either the stimulus or choice on each trial. We trained separate classifiers on every time bin in the training fold. We used the classifiers to predict stimulus or choice on each trial in the held-out test fold.

For both naive and balanced classifiers, we repeated the entire procedure five times, such that each trial was included in the test fold exactly once (and in the training fold the other four times). We averaged the classifier’s accuracy over each of the four held-out test sets (never including the training set) and reported this as the classifier’s overall cross-validated accuracy in Fig 4G.

Finally, for the right panel of Fig 4G, we zeroed out the spikes on all “sampling whisks” (defined above in the videographic methods). We also zeroed out spikes on the cycle preceding and the two cycles following each sampling whisk to ensure complete removal of whisk-locked stimulus information. This procedure removed phasic contact-evoked or whisk-evoked stimulus responses, but spared long-timescale persistent representations.

#### Neural encoding analysis (Fig 5, 6)

For this analysis, we began with the same features (contact count, etc.) from the behavioral analysis. Rather than aggregate within arbitrary time bins, we used the feature measurements on individual whisk cycles. We added some additional features that could affect neural firing: the amplitude (peak-to-trough angle) of each whisk and the set point (start angle) of each individual whisker at the beginning of each whisk.

We also added some additional trial-related features: current choice, previous choice, current outcome (rewarded or not), and previous outcome. Because the effect of these features could vary over the course of the trial, we used separate temporal indicator variables (Park et al., 2014). Specifically, we divided all whisks into 500 ms bins with respect to the response window opening. If the current choice was “left”, we marked the temporal indicator variable corresponding to left choices within that whisk’s bin as 1, and left all other variables as zero. We repeated this for each task variable.

Finally, we added two “nuisance features” for firing rate drift and cycle duration. For firing rate drift, we divided each session into 10 blocks and assessed the mean firing rate of each neuron within that block. We provided the logarithm of this value as a feature to the GLM. The timescale of each block was far too long (~several minutes) to contain any information about individual whisks, but it captured the baseline firing rate of the neuron, as well as any long-timescale variations, for example due to satiety. The second nuisance feature was the logarithm of the duration of each individual whisk cycle. This is because a whisk cycle that is twice as long should be expected to emit twice as many spikes, all else equal. The use of a logarithm in both cases accounts for the exponential link function in the GLM. Both of these nuisance features are highly predictive of neural firing by design and were important for fitting the data but were not analyzed further for scientific conclusions.

We fit the data using a GLM for Poisson data like spike counts (*i.e.*, with an exponential link function) using the ‘pyglmnet’ module (Jas et al., 2020). We used 5-fold cross-validation, ensuring that each trial was in the test set exactly once and evaluating the GLM on these held-out test sets only. We always used L2 regularization but we varied the strength of this regularization. We typically used the regularization value that optimized the model fit for that neuron, but when comparing coefficients across neurons (e.g., Fig 6) the same value of regularization was used for all neurons to ensure coefficients were on the same scale.

In order to obtain the null distributions of each coefficient and thereby significance, we also trained 40 additional GLMs for each neuron using permuted features. Specifically we permuted the rows but not the columns of the feature matrix, which maintains the correlation structure of the features but randomizes the mapping to neural responses. The distribution of each coefficient over permutations had a near-zero mean but a non-zero standard deviation. To assess significance of individual coefficients (e.g. Fig 5E-F) we divided the actual coefficient by the standard deviation of the coefficients obtained on the permutations to obtain the z-score of the coefficient. We then converted this into a two-tailed p-value by integrating the standard normal beyond this z-score. We validated that this approach controlled the false positive rate at α = 0.05 by including a spurious regressor that was drawn from a random distribution and ensuring that the random regressor was found significant no more than 5% of the time (indeed, that the resulting p-value distribution was uniform; data not shown).

To assess goodness-of-fit of any GLM, we took the log-likelihood of the data under the best fit and compared it to the log-likelihood of the data under a null model. The null model had access only to the “nuisance features” described above: baseline firing rate and whisk cycle duration. We subtracted the log-likelihood of the null from the log-likelihood of the fit model, and divided by the total number of whisks in that session in order to permit comparison across datasets of different duration. We used a logarithm of base 2 to permit presentation in “bits”. This is not an estimate of the information contained by the neural spike train, but rather an estimate of the change in the KL-divergence between [the true (unknown) distribution of the data and the distribution predicted by the model under consideration] versus [the same quantity, but replacing the model under consideration with the null model].

#### Analysis software

We used the Python packages ipython (Perez and Granger, 2007), pandas (McKinney, 2010), numpy (Van Der Walt et al., 2011), scipy (Virtanen et al., 2020), scikit-learn (Pedregosa et al., 2011), scikit-image (Van Der Walt et al., 2014), statsmodels, pyglmnet (Jas et al., 2020), and matplotlib (Hunter, 2007) to investigate, analyze, and present the data.

## Supplemental Information

**Supplemental Figure 1, related to Figure 1.**
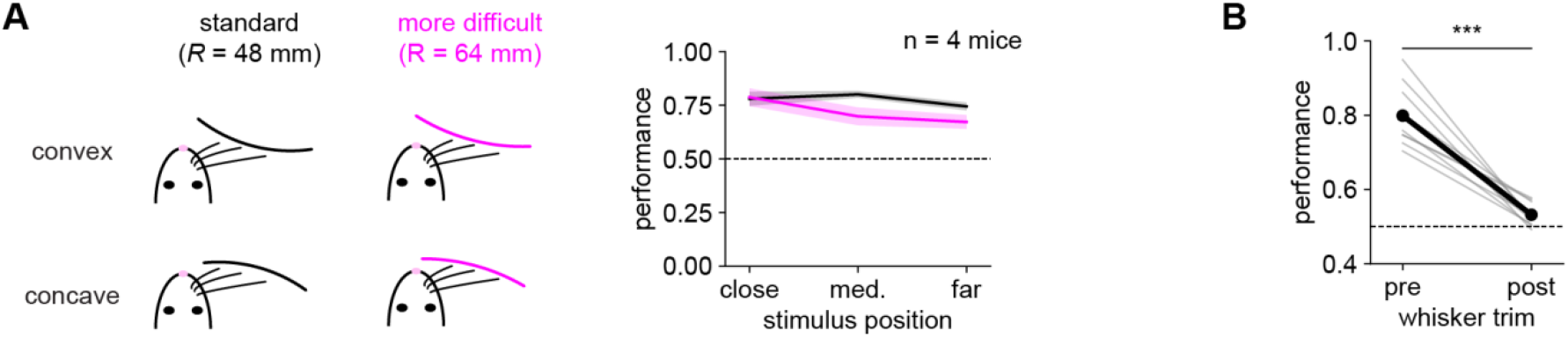
Generalization to more difficult stimuli and no-whisker controls. **A)** We trained a subset of mice on more difficult stimuli with a larger radius of curvature (pink). After some retraining, mice were able to perform well at these new stimuli, though at a slightly lower performance than for the original stimuli. Error bars: SEM over mice. **B)** Effect of trimming all whiskers on the performance of n = 10 mice performing the shape discrimination or detection tasks. “Pre”: average performance on the three sessions preceding whisker trim. “Post”: performance on the first session after whisker trim. Thin lines: individual mice. Thick line: average. Performance significantly decreased (paired t-test, p < .001) from 80.0% to 53.2%, near chance (50%, dashed line).

**Supplemental Figure 2, related to Figure 2.**
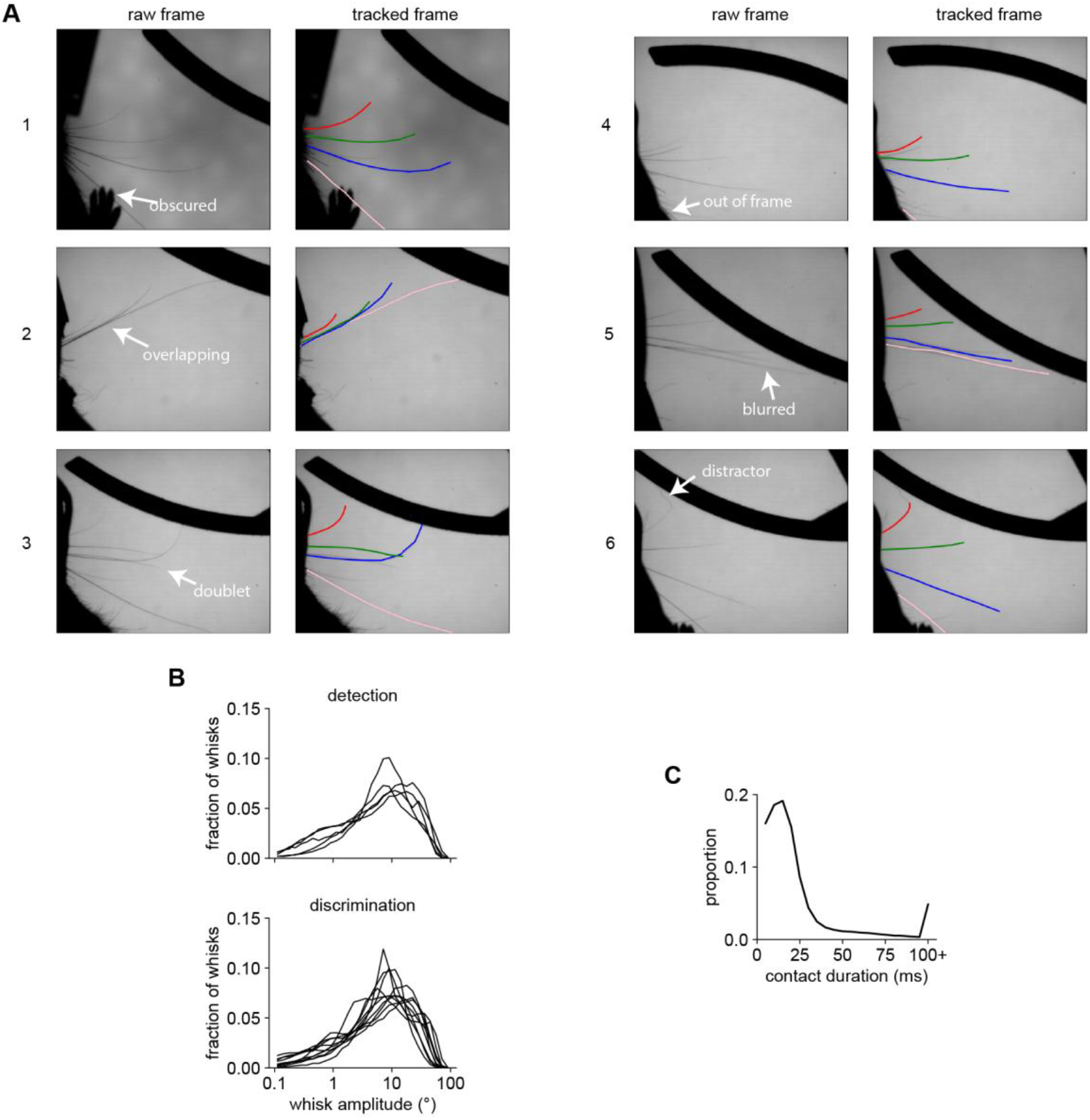
High-speed videography and whisking/contact statistics. **A)** Example frames demonstrating the quality of the whisker tracking. Within each pair of frames, the left frame is the raw frame (annotated with the region of interest) and the right frame shows the result of the whisker tracking algorithm. Performance was good (i.e., the correct whiskers were tracked throughout their extent) even when:

1. the whisker was obscured by a paw;
2. whiskers were nearly overlapping;
3. a “doublet” whisker emerged from the same follicle;
4. the whisker was nearly out of frame;
5. motion blurred the tips;
6. a similar-looking distractor hair was attached to the end of the whisker. **B)** Distribution of amplitudes of individual whisk cycles, with each line representing an individual mouse. Distributions were similar across mice, regardless of the task (top: detection; bottom: discrimination). **C)** Distribution of contact durations. Data were similar across tasks and therefore pooled. Most contacts were short but there was a long tail of longer contacts. Contact durations are quantized at the frame rate of 5 ms.

**Supplemental Figure 3, related to Figure 3.**
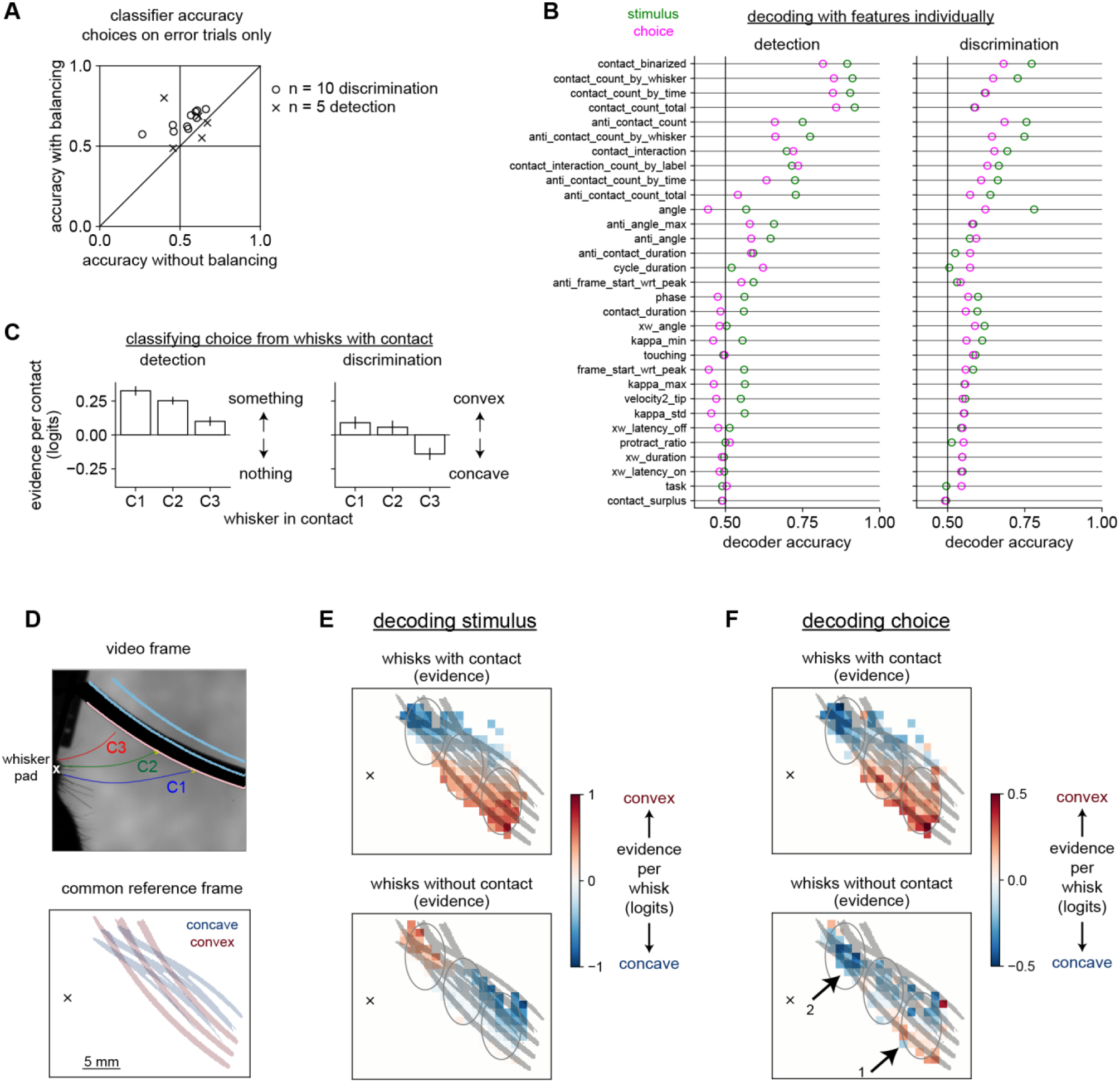
Behavioral decoding analysis. **A)** Accuracy of the behavioral decoders with and without trial balancing. The version described in the main manuscript uses balancing to equally weight correct and incorrect trials, and is plotted on the y-axis. We also trained a separate classifier that did not apply any balancing, but simply optimized its prediction of choice over all trials equally as in standard logistic regression; that accuracy is plotted on the x-axis. In both cases, the decoders had access to the full set of behavioral features. Here, we show the performance only on decoding choice on incorrect trials. Trial balancing improved the accuracy on this trial type in almost all cases and was always greater than 50%. On correct trials (data not shown), trial balancing slightly impaired performance, but was still quite high (*cf.* Fig 3D). **B)** (Related to Fig 3B) The performance of classifiers trained on every individual feature. See Supplemental Table 1 on the following page for descriptions of each features. **C)** (Related to Fig 3G) The weights used to predict choice from whisks with contact. The results were qualitatively similar to the weights used to predict stimulus (Fig 3G). During discrimination, C1 and C3 weights have opposite sign. **D,E)** Replotted from Fig 3I,K for comparison. **F)** The same data (top: whisks with contact; bottom: whisks without contact) now colored by the evidence they contain about choice, *i.e.* whether the mouse would report the shape to be concave (blue) or convex (red). The top panel is similar to the analogous panel in (E), indicating that mice correctly interpret whisks with contact in accordance with the ideal stimulus decoding. In the bottom panel, some whisks without contact are correctly interpreted by the mouse (that is, they look the same as in the analogous panel in (E)), but others are not. Specifically, in area 1, whisks without contact around the location of the closest convex shape indicate that the mouse would report convex. In area 2, whisks without contact around the location of the closest concave shape indicate that the mouse would report concave. This suggests that the mouse may be sampling these areas preferentially because it believes the corresponding shape is present.

**Supplemental Table 1.**
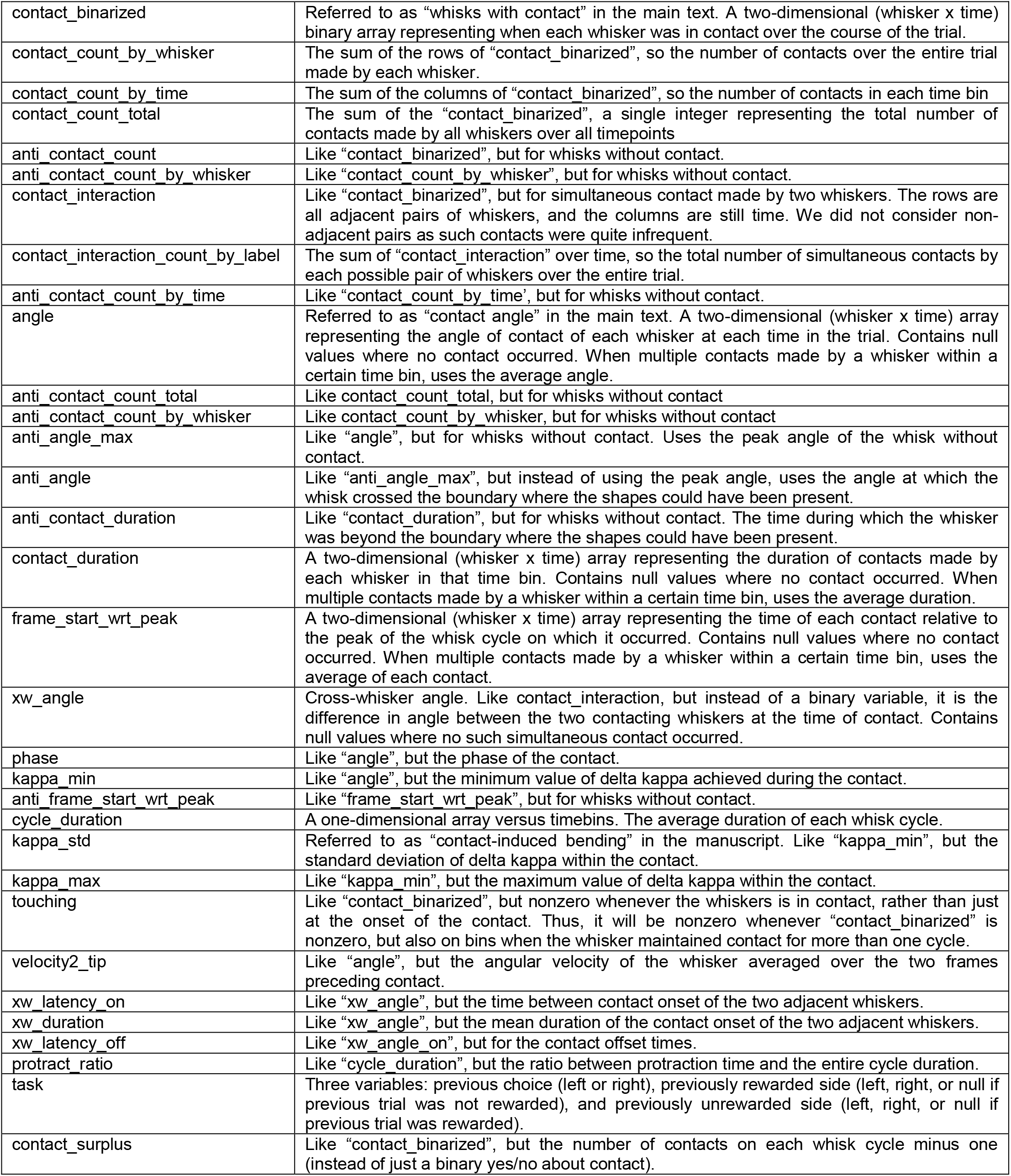
Defines the variables used in Supplemental Fig 3B, the entire list of features considered for inclusion in the behavioral decoders.

**Supplemental Figure 4, related to Figure 4.**
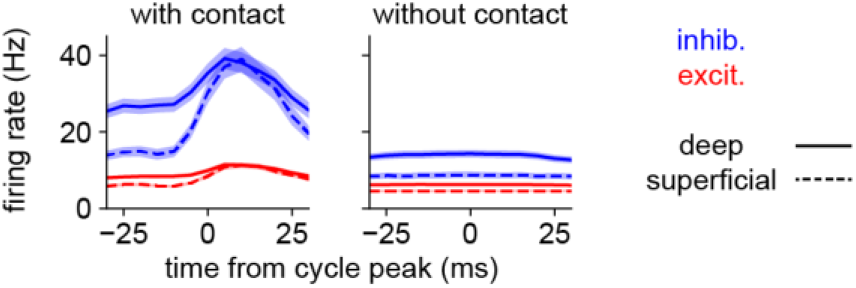
Neural responses to whisks with and without contact. The same data from Fig 4D, but plotted in Hz instead of normalized to baseline. The results are qualitatively similar, but the inhibitory cells (blue) show a much higher baseline firing rate than the excitatory cells (red). Error bars: SEM over neurons.

**Supplemental Figure 6, related to Figure 6.**
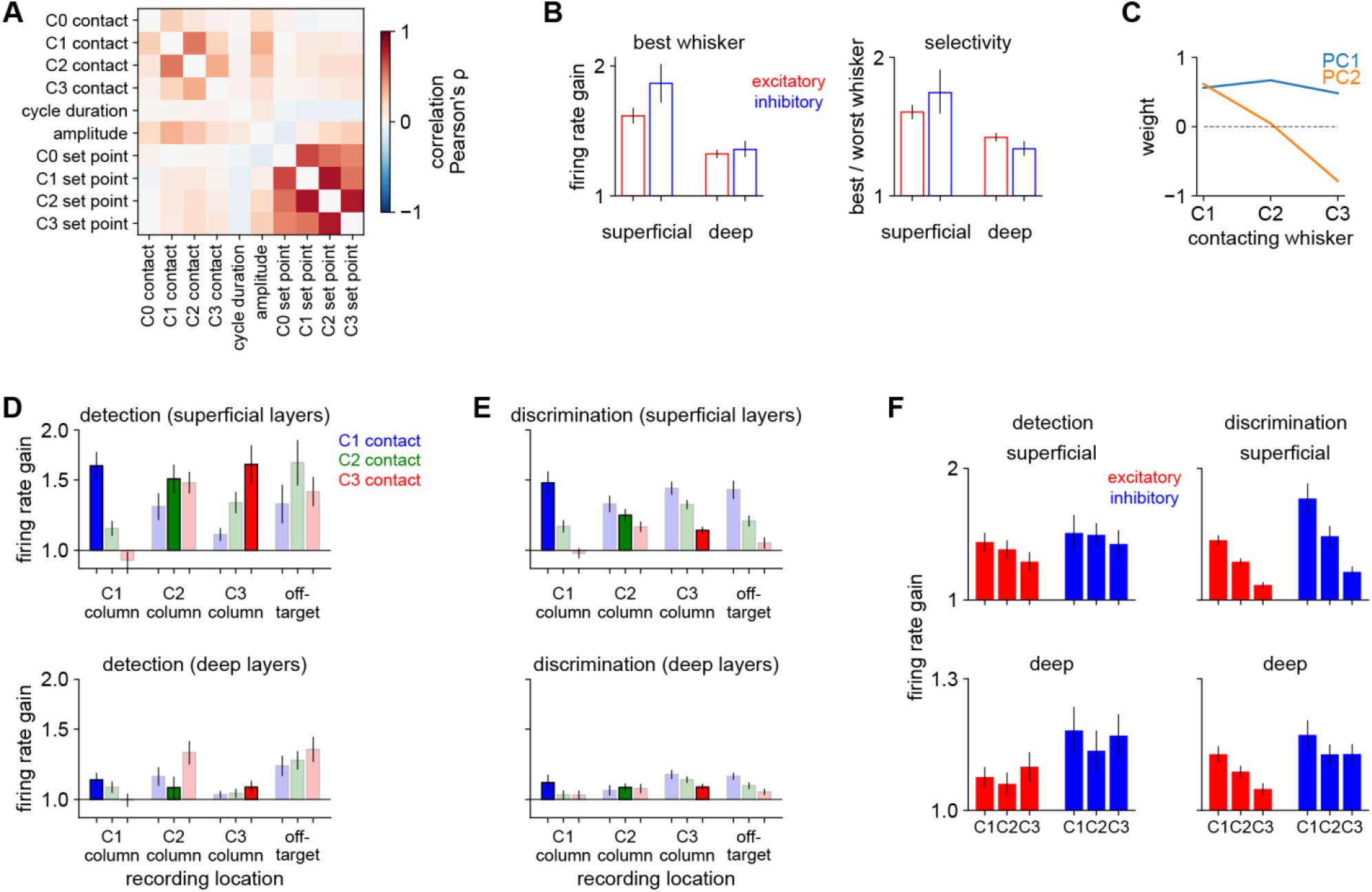
Task-specific contact responses and somatotopy. **A)** Feature correlation matrix. We concatenated the features over all sessions and calculated Pearson’s ρ between every pair of features. For clarity, task-related temporal indicator variables are not included here because they are orthogonal by design, and auto-correlation values along the diagonal are masked. **B)** *Left:* Similar to Fig 6E, but showing the response to the best whisker (*i.e.*, the whisker that evokes the strongest response in each neuron) instead of the average across C1-C3. The pattern is the same as in the main text: superficial neurons respond more strongly. *Right*: the selectivity of each neuron, parameterized as the response to the best whisker divided by the response to the worst whisker. Error bars: 95% bootstrapped confidence intervals. **C)** Principle component analysis (PCA) applied to the response of each neuron to contacts made by each whisker. The first PC (blue) is nearly equal on all whiskers whereas the second PC tracks the anterior-posterior position of whiskers C1-C3. Thus, the first PC captures overall response strength and the second PC captures topographic preference (C1>C3 or vice versa). These two PCs capture 63% and 28% of the variance in contact responses. **D)** The response to contacts by each whisker during shape detection, separately plotted by the cortical column in which they were recorded (C1, C2, C3, or “off-target”, meaning all other columns). Superficial neurons (top) show somatotopy, typically preferring their topographically aligned whisker (dark bars) over all others (light bars). Deep neurons (bottom) show weaker responses and less somatotopy. Error bars: SEM over neurons. **E)** Similar to (D), but for the shape discrimination task. The preference for C1 contacts (blue bars) dominates somatotopic responses (dark bars). **F)** Similar to (D) and (E), but pooling over recording locations and separately plotting by cell type (excitatory: red; inhibitory: blue). All populations prefer C1 contacts during discrimination (right). n = 235 neurons during detection and 675 neurons during discrimination. We excluded sessions in which too few contacts were made by any whisker (C1-C3) to estimate the response. Logarithmic y-axis in B (left) and D-F.

**Supplemental Video 1.**
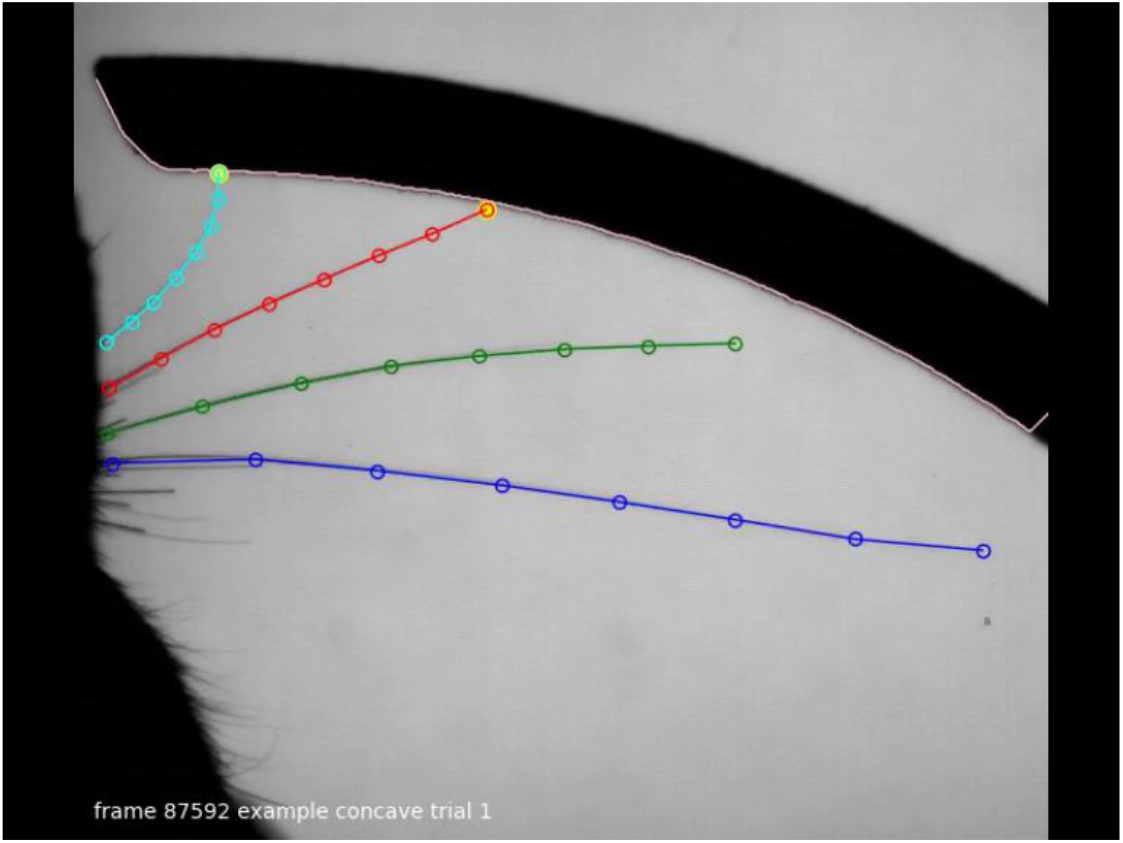
This video demonstrates some example trials of the shape discrimination behavior. A single frame is shown here for illustration. The whisker tracking is overlaid (C0, blue; C1, green; C2, red; C3, cyan; circles indicate individually tracked “joints”). Yellow circles appear when any tip contacts the edge of the shape (pink line). Four example correct trials (two concave and two convex) are included. The playback speed is 0.15x real-time.

**Supplemental Video 2.**
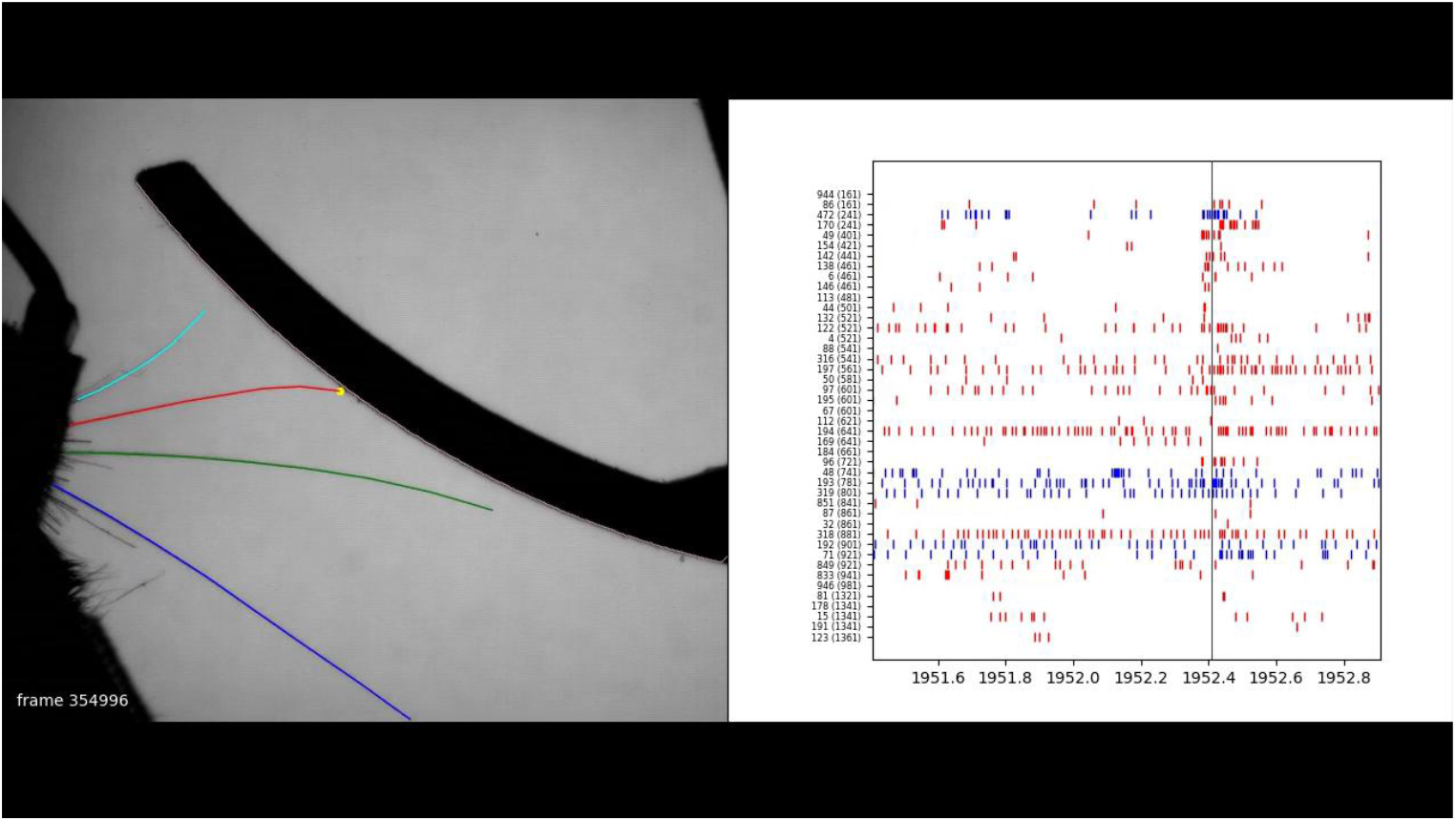
This video demonstrates simultaneously recorded neural activity and behavior. A single frame is shown here for illustration. The left half of the frame shows the behavior, as in Supplemental Video 1 (excluding the tracked joints for clarity). The right half of the frame shows the neural spiking, synchronized with the behavior. Individual neurons (red: excitatory; blue: inhibitory) are plotted as separate rows. Neurons are sorted from superficial (top) to deep (bottom). The small text labels next to each row indicate the unit number and, in parentheses, the depth in microns from the cortical surface. An audio track plays the spikes from single example neurons, as annotated throughout the video. As in Supplemental Video 1, the playback speed is 0.15x real-time.

## Notes

### Competing Interest Statement

The authors have declared no competing interest.

## References

Adesnik, H., and Naka, A. (2018). Cracking the Function of Layers in the Sensory Cortex. Neuron 100, 1028–1043.

Ahissar, E., and Assa, E. (2016). Perception as a closed-loop convergence process. Elife 5, 1–26.

Akrami, A., Kopec, C.D., Diamond, M.E., and Brody, C.D. (2018). Posterior parietal cortex represents sensory history and mediates its effects on behaviour. Nature 554, 368–372.

Anjum, F., Turni, H., Mulder, P.G.H., van der Burg, J., and Brecht, M. (2006). Tactile guidance of prey capture in Etruscan shrews. Proc. Natl. Acad. Sci. U. S. A. 103, 16544–16549.

Ayaz, A., Stäuble, A., Hamada, M., Wulf, M.A., Saleem, A.B., and Helmchen, F. (2019). Layer-specific integration of locomotion and sensory information in mouse barrel cortex. Nat. Commun. 10.

Bale, M.R., and Maravall, M. (2018). Organization of sensory feature selectivity in the whisker system. Neuroscience 368, 70–80.

Bandyopadhyay, S., Shamma, S.A., and Kanold, P.O. (2010). Dichotomy of functional organization in the mouse auditory cortex. Nat. Neurosci. 13, 361–368.

Benison, A.M., Ard, T.D., Crosby, A.M., and Barth, D.S. (2006). Temporal patterns of field potentials in vibrissa/barrel cortex reveal stimulus orientation and shape. J. Neurophysiol. 95, 2242–2251.

Birdwell, J.A., Solomon, J.H., Thajchayapong, M., Taylor, M.A., Cheely, M., Towal, R.B., Conradt, J., and Hartmann, M.J.Z. (2007). Biomechanical models for radial distance determination by the rat vibrissal system. J. Neurophysiol. 98, 2439–2455.

Bortone, D.S., Olsen, S.R., and Scanziani, M. (2014). Translaminar inhibitory cells recruited by layer 6 corticothalamic neurons suppress visual cortex. Neuron 82, 474–485.

Brecht, M. (2017). The Body Model Theory of Somatosensory Cortex. Neuron 94, 985–992.

Brecht, M., Preilowski, B., and Merzenich, M.M. (1997). Functional architecture of the mystacial vibrissae. Behav. Brain Res. 84, 81–97.

Brecht, M., Roth, A., and Sakmann, B. (2003). Dynamic receptive fields of reconstructed pyramidal cells in layers 3 and 2 of rat somatosensory barrel cortex. J. Physiol. 553, 243–265.

Brown, J., Oldenburg, I.A., Telian, G.I., Griffin, S., Voges, M., Jain, V., and Adesnik, H. (2020). Spatial integration during active tactile sensation drives elementary shape perception. BioRxiv.

Brumberg, J.C., Pinto, D.J., and Simons, D.J. (1996). Spatial gradients and inhibitory summation in the rat whisker barrel system. J. Neurophysiol. 76, 130–140.

Bruno, R.M., and Sakmann, B. (2006). Cortex is driven by weak but synchronously active thalamocortical synapses. Science (80-.). 312, 1622–1627.

Bruno, R.M., and Simons, D.J. (2002). Feedforward mechanisms of excitatory and inhibitory cortical receptive fields. J. Neurosci. 22, 10966–10975.

Bush, N.E., Solla, S.A., and Hartmann, M.J. (2016). Whisking mechanics and active sensing. Curr. Opin. Neurobiol. 40, 178–188.

Campagner, D., Evans, M.H., Bale, M.R., Erskine, A., and Petersen, R.S. (2016). Prediction of primary somatosensory neuron activity during active tactile exploration. Elife 5, 1–18.

Campagner, D., Evans, M.H., Chlebikova, K., Colins-Rodriguez, A., Loft, M.S.E., Fox, S., Pettifer, D., Humphries, M.D., Svoboda, K., and Petersen, R.S. (2019). Prediction of choice from competing mechanosensory and choice-memory cues during active tactile decision making. J. Neurosci. 39, 3921–3933.

Carcea, I., Insanally, M.N., and Froemke, R.C. (2017). Dynamics of auditory cortical activity during behavioural engagement and auditory perception. Nat. Commun. 8, 1–12.

Cardona, A., Saalfeld, S., Schindelin, J., Arganda-Carreras, I., Preibisch, S., Longair, M., Tomancak, P., Hartenstein, V., and Douglas, R.J. (2012). TrakEM2 software for neural circuit reconstruction. PLoS One 7.

Carvell, G.E., and Simons, D.J. (1990). Biometric analyses of vibrissal tactile discrimination in the rat. J. Neurosci. 10, 2638–2648.

Carvell, G.E., and Simons, D.J. (1995). Task- and subject-related differences in sensorimotor behavior during active touch. Somatosens. Mot. Res. 12, 1–9.

Celikel, T., and Sakmann, B. (2007). Sensory integration across space and in time for decision making in the somatosensory system of rodents. Proc. Natl. Acad. Sci. U. S. A. 104, 1395–1400.

Chapman, C.E., and Ageranioti-Bélanger, S.A. (1991). Discharge properties of neurones in the hand area of primary somatosensory cortex in monkeys in relation to the performance of an active tactile discrimination task - I. Areas 3b and 1. Exp. Brain Res. 87, 319–339.

Chen, J.L., Carta, S., Soldado-Magraner, J., Schneider, B.L., and Helmchen, F. (2013). Behaviour-dependent recruitment of long-range projection neurons in somatosensory cortex. Nature 499, 336–340.

Cheung, J., Maire, P., Kim, J., Sy, J., Cheung, J., Maire, P., Kim, J., Sy, J., and Hires, S.A. (2019). The Sensorimotor Basis of Whisker-Guided Anteroposterior Object Localization in Head-Fixed Mice. Curr. Biol. 1–12.

Clack, N.G., O’Connor, D.H., Huber, D., Petreanu, L., Hires, A., Peron, S., Svoboda, K., and Myers, E.W. (2012). Automated tracking of whiskers in videos of head fixed rodents. PLoS Comput. Biol. 8, e1002591.

Clancy, K.B., Schnepel, P., Rao, A.T., and Feldman, D.E. (2015). Structure of a Single Whisker Representation in Layer 2 of Mouse Somatosensory Cortex. J. Neurosci. 35, 3946–3958.

Connor, C.E., Brincat, S.L., and Pasupathy, A. (2007). Transformation of shape information in the ventral pathway. Curr. Opin. Neurobiol. 17, 140–147.

Constantinople, C.M., and Bruno, R.M. (2013). Deep cortical layers are activated directly by thalamus. Science 340, 1591–1594.

Cruikshank, S.J., Lewis, T.J., and Connors, B.W. (2007). Synaptic basis for intense thalamocortical activation of feedforward inhibitory cells in neocortex. Nat. Neurosci. 10, 462–468.

Curtis, J.C., and Kleinfeld, D. (2009). Phase-to-rate transformations encode touch in cortical neurons of a scanning sensorimotor system. Nat. Neurosci. 12, 492–501.

Dabney, W., Kurth-Nelson, Z., Uchida, N., Starkweather, C.K., Hassabis, D., Munos, R., and Botvinick, M. (2020). A distributional code for value in dopamine-based reinforcement learning. Nature 577, 671–675.

Davidson, P.W. (1972). Haptic judgments of curvature by blind and sighted humans. J. Exp. Psychol. 93, 43–55.

Diamond, M.E. (2010). Texture sensation through the fingertips and the whiskers. Curr. Opin. Neurobiol. 20, 319–327.

Diamond, M.E., von Heimendahl, M., Knutsen, P.M., Kleinfeld, D., and Ahissar, E. (2008). “Where” and “what” in the whisker sensorimotor system. Nat. Rev. Neurosci. 9, 601–612.

Dominiak, S.E., Nashaat, M.A., Sehara, K., Oraby, H., Larkum, M.E., and Sachdev, R.N.S. (2019). Whisking Asymmetry Signals Motor Preparation and the Behavioral State of Mice. J. Neurosci. 39, 9818–9830.

Douglas, R.J., and Martin, K.A.C. (2004). Neuronal circuits of the neocortex. Annu. Rev. Neurosci. 27, 419–451.

Drew, P.J., and Feldman, D.E. (2007). Representation of moving wavefronts of whisker deflection in rat somatosensory cortex. J. Neurophysiol. 98, 1566–1580.

Ego-Stengel, V., Mello e Souza, T., Jacob, V., and Shulz, D.E. (2005). Spatiotemporal characteristics of neuronal sensory integration in the barrel cortex of the rat. J. Neurophysiol. 93, 1450–1467.

von der Emde (2010). 3-Dimensional scene perception during active electrolocation in a weakly electric pulse fish. Front. Behav. Neurosci. 4, 1–13.

Estebanez, L., Férézou, I., Ego-Stengel, V., and Shulz, D.E. (2018). Representation of tactile scenes in the rodent barrel cortex. Neuroscience 368, 81–94.

Fassihi, A., Zuo, Y., and Diamond, M.E. (2020). Making sense of sensory evidence in the rat whisker system. Curr. Opin. Neurobiol. 60, 76–83.

Fee, M.S., Mitra, P.P., and Kleinfeld, D. (1997). Central versus peripheral determinants of patterned spike activity in rat vibrissa cortex during whisking. J. Neurophysiol. 78, 1144–1149.

Frandolig, J.E., Matney, C.J., Lee, K., Kim, J., Chevée, M., Kim, S.J., Bickert, A.A., and Brown, S.P. (2019). The Synaptic Organization of Layer 6 Circuits Reveals Inhibition as a Major Output of a Neocortical Sublamina. Cell Rep. 28, 3131–3143.e5.

Gamzu, E., and Ahissar, E. (2001). Importance of temporal cues for tactile spatial-frequency discrimination. J Neurosci 21, 7416–7427.

Gibson, J.J. (1962). Observations on active touch. Psychol. Rev. 69, 477–491.

Goodman, J.M., Tabot, G.A., Lee, A.S., Suresh, A.K., Rajan, A.T., Hatsopoulos, N.G., and Bensmaia, S. (2019). Postural Representations of the Hand in the Primate Sensorimotor Cortex. Neuron 104, 1000–1009.e7.

Gouwens, N.W., Sorensen, S.A., Berg, J., Lee, C., Jarsky, T., Ting, J., Sunkin, S.M., Feng, D., Anastassiou, C.A., Barkan, E., et al. (2019). Classification of electrophysiological and morphological neuron types in the mouse visual cortex. Nat. Neurosci. 22, 1182–1195.

Grant, R.A., Mitchinson, B., Fox, C.W., and Prescott, T.J. (2009). Active Touch Sensing in the Rat: Anticipatory and Regulatory Control of Whisker Movements During Surface Exploration. J. Neurophysiol. 101, 862–874.

Grant, R.A., Breakell, V., and Prescott, T.J. (2018). Whisker touch sensing guides locomotion in small, quadrupedal mammals. Proc. R. Soc. B Biol. Sci. 285.

Guo, Z. V., Hires, S.A., Li, N., O’Connor, D.H., Komiyama, T., Ophir, E., Huber, D., Bonardi, C., Morandell, K., Gutnisky, D., et al. (2014). Procedures for Behavioral Experiments in Head-Fixed Mice. PLoS One 9, e88678.

Gustafson, J.W., and Felbain-Keramidas, S.L. (1977). Behavioral and neural approaches to the function of the mystacial vibrissae. Psychol. Bull. 84, 477–488.

Harvey, M.A., Bermejo, R., and Zeigler, H.P. (2001). Discriminative whisking in the head-fixed rat: optoelectronic monitoring during tactile detection and discrimination tasks. Somatosens. Mot. Res. 18, 211–222.

Hattori, R., Danskin, B., Babic, Z., Mlynaryk, N., and Komiyama, T. (2019). Area-Specificity and Plasticity of History-Dependent Value Coding During Learning. Cell 177, 1858–1872.e15.

He, K., Zhang, X., Ren, S., and Sun, J. (2015). Deep residual learning for image recognition. ArXiv.

Hill, D.N., Curtis, J.C., Moore, J.D., and Kleinfeld, D. (2011). Primary motor cortex reports efferent control of vibrissa motion on multiple timescales. Neuron 72, 344–356.

Hires, S.A., Pammer, L., Svoboda, K., and Golomb, D. (2013). Tapered whiskers are required for active tactile sensation. Elife 2, e01350.

Hires, S.A., Gutnisky, D.A., Yu, J., O’Connor, D.H., and Svoboda, K. (2015). Low-noise encoding of active touch by layer 4 in the somatosensory cortex. Elife 4, 1–18.

Hobbs, J.A., Towal, R.B., and Hartmann, M.J.Z. (2016a). Spatiotemporal Patterns of Contact Across the Rat Vibrissal Array During Exploratory Behavior. Front. Behav. Neurosci. 9, 1–18.

Hobbs, J.A., Towal, R.B., and Hartmann, M.J.Z. (2016b). Evidence for Functional Groupings of Vibrissae across the Rodent Mystacial Pad. PLoS Comput. Biol. 12, 1–36.

Hong, Y.K., Lacefield, C.O., Rodgers, C.C., and Bruno, R.M. (2018). Sensation, movement and learning in the absence of barrel cortex. Nature 561, 542–546.

Hooks, B.M., Hires, S.A., Zhang, Y.-X., Huber, D., Petreanu, L., Svoboda, K., and Shepherd, G.M.G. (2011). Laminar analysis of excitatory local circuits in vibrissal motor and sensory cortical areas. PLoS Biol. 9, e1000572.

Huber, D., Gutnisky, D.A., Peron, S., O’Connor, D.H., Wiegert, J.S., Tian, L., Oertner, T.G., Looger, L.L., and Svoboda, K. (2012). Multiple dynamic representations in the motor cortex during sensorimotor learning. Nature 484, 473–478.

Hunter, J.D. (2007). Matplotlib: a 2d graphics environment. Comput. Sci. Eng. 9, 90–95.

Hutson, K.A., and Masterton, R.B. (1986). The sensory contribution of a single vibrissa’s cortical barrel. J. Neurophysiol. 56, 1196–1223.

Insafutdinov, E., Pishchulin, L., Andres, B., Andriluka, M., and Schiele, B. (2016). DeeperCut: A Deeper, Stronger, and Faster Multi-Person Pose Estimation Model. Eur. Conf. Comput. Vis. 34–50.

Isett, B.R., Feasel, S.H., Lane, M.A., and Feldman, D.E. (2018). Slip-Based Coding of Local Shape and Texture in Mouse S1. Neuron 1–16.

Jacob, V., Le Cam, J., Ego-Stengel, V., and Shulz, D.E. (2008). Emergent properties of tactile scenes selectively activate barrel cortex neurons. Neuron 60, 1112–1125.

Jadhav, S.P., Wolfe, J., and Feldman, D.E. (2009). Sparse temporal coding of elementary tactile features during active whisker sensation. Nat. Neurosci. 12, 792–800.

Jang, Y., Wixted, J.T., and Huber, D.E. (2009). Testing Signal-Detection Models of Yes/No and Two-Alternative Forced-Choice Recognition Memory. J. Exp. Psychol. Gen. 138, 291–306.

Jas, M., Achakulvisut, T., Idrizović, A., Acuna, D., Antalek, M., Marques, V., Odland, T., Garg, R., Agrawal, M., Umegaki, Y., et al. (2020). Pyglmnet: Python implementation of elastic-net regularized generalized linear models. J. Open Source Softw. 5, 1959.

Johansson, R.S., and Flanagan, J.R. (2009). Coding and use of tactile signals from the fingertips in object manipulation tasks. Nat. Rev. Neurosci. 10, 345–359.

Keller, G.B., and Mrsic-Flogel, T.D. (2018). Predictive Processing: A Canonical Cortical Computation. Neuron 100, 424–435.

Keller, G.B., Bonhoeffer, T., and Hübener, M. (2012). Sensorimotor mismatch signals in primary visual cortex of the behaving mouse. Neuron 74, 809–815.

Kinnischtzke, A.K., Simons, D.J., and Fanselow, E.E. (2014). Motor cortex broadly engages excitatory and inhibitory neurons in somatosensory barrel cortex. Cereb. Cortex 24, 2237–2248.

Klatzky, R.L., and Lederman, S.J. (2011). Haptic object perception: Spatial dimensionality and relation to vision. Philos. Trans. R. Soc. B Biol. Sci. 366, 3097–3105.

Knutsen, P.M., Pietr, M., and Ahissar, E. (2006). Haptic object localization in the vibrissal system: behavior and performance. J. Neurosci. 26, 8451–8464.

De Kock, C.P.J., Bruno, R.M., Spors, H., and Sakmann, B. (2007). Layer- and cell-type-specific suprathreshold stimulus representation in rat primary somatosensory cortex. J. Physiol. 581, 139–154.

Krakauer, J.W., Ghazanfar, A.A., Gomez-Marin, A., Maciver, M.A., and Poeppel, D. (2017). Neuroscience Needs Behavior: Correcting a Reductionist Bias. Neuron 93, 480–490.

Krupa, D.J., Matell, M.S., Brisben, A.J., Oliveira, L.M., and Nicolelis, M.A. (2001). Behavioral properties of the trigeminal somatosensory system in rats performing whisker-dependent tactile discriminations. J. Neurosci. 21, 5752–5763.

Krupa, D.J., Wiest, M.C., Shuler, M.G., Laubach, M., and Nicolelis, M.A.L. (2004). Layer-specific somatosensory cortical activation during active tactile discrimination. Science 304, 1989–1992.

Laboy-Juárez, K.J., Langberg, T., Ahn, S., and Feldman, D.E. (2019). Elementary motion sequence detectors in whisker somatosensory cortex. Nat. Neurosci. 22, 1438–1449.

Lacefield, C.O., Pnevmatikakis, E.A., Paninski, L., and Bruno, R.M. (2019). Reinforcement Learning Recruits Somata and Apical Dendrites across Layers of Primary Sensory Cortex. Cell Rep. 26, 2000–2008.e2.

Lavzin, M., Levy, S., Benisty, H., Dubin, U., Brosh, Z., Aeed, F., Mensh, B.D., Schiller, Y., Meir, R., Barak, O., et al. (2020). Cell-type specific outcome representation in primary motor cortex. BioRxiv.

Lederman, S.J., and Klatzky, R.L. (1987). Hand movements: a window into haptic object recognition. Cogn. Psychol. 19, 342–368.

Lyon, L., Saksida, L.M., and Bussey, T.J. (2012). Spontaneous object recognition and its relevance to schizophrenia: a review of findings from pharmacological, genetic, lesion and developmental rodent models. Psychopharmacology (Berl). 220, 647–672.

Marr, D.C., and Poggio, T. (1976). From understanding computation to understanding neural circuitry. MIT AI Memos 357, 1–22.

Mathis, A., Mamidanna, P., Cury, K.M., Abe, T., Murthy, V.N., Mathis, M.W., and Bethge, M. (2018). DeepLabCut: markerless pose estimation of user-defined body parts with deep learning. Nat. Neurosci. 21, 1281–1289.

McGinley, M.J., David, S.V., and McCormick, D.A. (2015). Cortical Membrane Potential Signature of Optimal States for Sensory Signal Detection. Neuron 1–14.

McKinney, W. (2010). Data structures for statistical computing in Python. Proc. 9th Python Sci. Conf.

Mehta, S.B., Whitmer, D., Figueroa, R., Williams, B.A., and Kleinfeld, D. (2007). Active spatial perception in the vibrissa scanning sensorimotor system. PLoS Biol. 5, e15.

Mitchinson, B., Martin, C.J., Grant, R.A., and Prescott, T.J. (2007). Feedback control in active sensing: Rat exploratory whisking is modulated by environmental contact. Proc. R. Soc. B Biol. Sci. 274, 1035–1041.

Mohan, H., de Haan, R., Broersen, R., Pieneman, A.W., Helmchen, F., Staiger, J.F., Mansvelder, H.D., and de Kock, C.P.J. (2019). Functional Architecture and Encoding of Tactile Sensorimotor Behavior in Rat Posterior Parietal Cortex. J. Neurosci. 39, 7332–7343.

Moore, J.D., Mercer Lindsay, N., Deschênes, M., and Kleinfeld, D. (2015). Vibrissa Self-Motion and Touch Are Reliably Encoded along the Same Somatosensory Pathway from Brainstem through Thalamus. PLoS Biol. 13, e1002253.

Muñoz, W., Tremblay, R., Levenstein, D., and Rudy, B. (2017). Layer-specific modulation of neocortical dendritic inhibition during active wakefulness. Science (80-.). 959, 954–959.

Musall, S., Kaufman, M.T., Juavinett, A.L., Gluf, S., and Churchland, A.K. (2019). Single-trial neural dynamics are dominated by richly varied movements. Nat. Neurosci. 22, 1677–1686.

Nandy, A., Sharpee, T., Reynolds, J., and Mitchell, J. (2013). The Fine Structure of Shape Tuning in Area V4. Neuron 78, 1102–1115.

Nogueira, R., Abolafia, J.M., Drugowitsch, J., Balaguer-Ballester, E., Sanchez-Vives, M. V., and Moreno-Bote, R. (2017). Lateral orbitofrontal cortex anticipates choices and integrates prior with current information. Nat. Commun. 8.

Nogueira, R., Peltier, N.E., Anzai, A., DeAngelis, G.C., Martínez-Trujillo, J., and Moreno-Bote, R. (2020). The effects of population tuning and trial-by-trial variability on information encoding and behavior. J. Neurosci. 40, 1066–1083.

O’Connor, D.H., Clack, N.G., Huber, D., Komiyama, T., Myers, E.W., and Svoboda, K. (2010a). Vibrissa-based object localization in head-fixed mice. J. Neurosci. 30, 1947–1967.

O’Connor, D.H., Peron, S.P., Huber, D., and Svoboda, K. (2010b). Neural activity in barrel cortex underlying vibrissa-based object localization in mice. Neuron 67, 1048–1061.

O’Connor, D.H., Hires, S.A., Guo, Z. V, Li, N., Yu, J., Sun, Q.-Q., Huber, D., and Svoboda, K. (2013). Neural coding during active somatosensation revealed using illusory touch. Nat. Neurosci. 16, 958–965.

Pachitariu, M., Steinmetz, N., Kadir, S., Carandini, M., and Harris, K.D. (2016). Kilosort: realtime spike-sorting for extracellular electrophysiology with hundreds of channels. BioRxiv 061481.

Pammer, L., O’Connor, D.H., Hires, S.A., Clack, N.G., Huber, D., Myers, E.W., and Svoboda, K. (2013). The mechanical variables underlying object localization along the axis of the whisker. J. Neurosci. 33, 6726–6741.

Park, I.M., Meister, M.L.R., Huk, A.C., and Pillow, J.W. (2014). Encoding and decoding in parietal cortex during sensorimotor decision-making. Nat. Neurosci. 17, 1395–1403.

Pedregosa, F., Varoquaux, G., Gramfort, A., Michel, V., Thirion, B., Grisel, O., Blondel, M., Prettenhofer, P., Weiss, R., Dubourg, V., et al. (2011). Scikit-learn. J. Mach. Learn. Res. 12, 2825–2830.

Perez, F., and Granger, B.E. (2007). IPython: A System for Interactive Scientific Computing. Comput. Sci. Eng. 9.

Peron, S.P., Freeman, J., Iyer, V., Guo, C., and Svoboda, K. (2015). A Cellular Resolution Map of Barrel Cortex Activity during Tactile Behavior. Neuron 1–17.

Petersen, R.S., Rodriguez, A.C., Evans, M.H., Campagner, D., and Loft, M.S.E. (2020). A system for tracking whisker kinematics and whisker shape in three dimensions. PLoS Comput. Biol. 16, 1–24.

Petreanu, L., Gutnisky, D.A., Huber, D., Xu, N., O’Connor, D.H., Tian, L., Looger, L., and Svoboda, K. (2012). Activity in motor-sensory projections reveals distributed coding in somatosensation. Nature 489, 299–303.

Pishchulin, L., Insafutdinov, E., Tang, S., Andres, B., Andriluka, M., Gehler, P., and Schiele, B. (2015). DeepCut: Joint Subset Partition and Labeling for Multi Person Pose Estimation.

Pluta, S.R., Lyall, E.H., Telian, G.I., Ryapolova-Webb, E., and Adesnik, H. (2017). Surround Integration Organizes a Spatial Map during Active Sensation. Neuron 94, 1220–1233.e5.

Pluta, S.R., Telian, G.I., Naka, A., and Adesnik, H. (2019). Superficial layers suppress the deep layers to fine-tune cortical coding. J. Neurosci. 39, 2052–2064.

Polley, D.B., Rickert, J.L., and Frostig, R.D. (2005). Whisker-based discrimination of object orientation determined with a rapid training paradigm. Neurobiol. Learn. Mem. 83, 134–142.

Quist, B.W., Seghete, V., Huet, L.A., Murphey, T.D., and Hartmann, M.J.Z. (2014). Modeling Forces and Moments at the Base of a Rat Vibrissa during Noncontact Whisking and Whisking against an Object. J. Neurosci. 34, 9828–9844.

Ramalingam, N., McManus, J.N.J., Li, W., and Gilbert, C.D. (2013). Top-down modulation of lateral interactions in visual cortex. J. Neurosci. 33, 1773–1789.

Ramirez, A., Pnevmatikakis, E.A., Merel, J., Paninski, L., Miller, K.D., and Bruno, R.M. (2014). Spatiotemporal receptive fields of barrel cortex revealed by reverse correlation of synaptic input. Nat. Neurosci. 17, 866–875.

Ranganathan, G.N., Apostolides, P.F., Harnett, M.T., Xu, N.L., Druckmann, S., and Magee, J.C. (2018). Active dendritic integration and mixed neocortical network representations during an adaptive sensing behavior. Nat. Neurosci. 21, 1583–1590.

Rigotti, M., Barak, O., Warden, M.R., Wang, X.-J., Daw, N.D., Miller, E.K., and Fusi, S. (2013). The importance of mixed selectivity in complex cognitive tasks. Nature 497, 585–590.

Ritt, J.T., Andermann, M.L., and Moore, C.I. (2008). Embodied information processing: vibrissa mechanics and texture features shape micromotions in actively sensing rats. Neuron 57, 599–613.

Robertson, C.E., and Baron-Cohen, S. (2017). Sensory perception in autism. Nat. Rev. Neurosci. 18, 671–684.

Rossant, C., Kadir, S.N., Goodman, D.F.M., Schulman, J., Hunter, M.L.D., Saleem, A.B., Grosmark, A., Belluscio, M., Denfield, G.H., Ecker, A.S., et al. (2016). Spike sorting for large, dense electrode arrays. Nat. Neurosci. 19, 634–641.

Rothschild, G., Nelken, I., and Mizrahi, A. (2010). Functional organization and population dynamics in the mouse primary auditory cortex. Nat. Neurosci. 13, 353–360.

Sachdev, R.N.S., Sellien, H., and Ebner, F. (2001). Temporal organization of multi-whisker contact in rats. Somatosens. Mot. Res. 18, 91–100.

Schneider, D.M., Sundararajan, J., and Mooney, R. (2018). A cortical filter that learns to suppress the acoustic consequences of movement. Nature 561, 391–395.

Schulman, A.J., and Mitchell, R.R. (1966). Operating Characteristics from Yes-No and Forced-Choice Procedures. J. Acoust. Soc. Am. 40, 473–477.

Schwarz, C. (2016). The Slip Hypothesis: Tactile Perception and its Neuronal Bases. Trends Neurosci. 39, 449–462.

Severson, K.S., Xu, D., Yang, H., and O’Connor, D.H. (2019). Coding of whisker motion across the mouse face. Elife 8, 1–23.

Sharpee, T.O. (2013). Computational Identification of Receptive Fields. Annu. Rev. Neurosci. 36, 103–120.

Sippy, T., Lapray, D., Crochet, S., and Petersen, C.C.H. (2015). Cell-Type-Specific Sensorimotor Processing in Striatal Projection Neurons during Goal-Directed Behavior. Neuron 1–8.

Steinmetz, N.A., and Moore, T. (2010). Changes in the response rate and response variability of area V4 neurons during the preparation of saccadic eye movements. J. Neurophysiol. 103, 1171–1178.

Stringer, C., Pachitariu, M., Steinmetz, N., Reddy, C.B., Carandini, M., and Harris, K.D. (2019). Spontaneous behaviors drive multidimensional, brainwide activity. Science (80-.). 364.

Stüttgen, M.C., and Schwarz, C. (2018). Barrel cortex: What is it good for? Neuroscience 368, 3–16.

Thakur, P.H., Bastian, A.J., and Hsiao, S.S. (2008). Multidigit movement synergies of the human hand in an unconstrained haptic exploration task. J. Neurosci. 28, 1271–1281.

Thakur, P.H., Fitzgerald, P.J., and Hsiao, S.S. (2012). Second-order receptive fields reveal multidigit interactions in area 3b of the macaque monkey. J. Neurophysiol. 108, 243–262.

Tsutsui, K.I., Grabenhorst, F., Kobayashi, S., and Schultz, W. (2016). A dynamic code for economic object valuation in prefrontal cortex neurons. Nat. Commun. 7.

Vilarchao, M.E., Estebanez, L., Shulz, D.E., and Férézou, I. (2018). Supra-barrel Distribution of Directional Tuning for Global Motion in the Mouse Somatosensory Cortex. Cell Rep. 22, 3534–3547.

Vinck, M., Batista-Brito, R., Knoblich, U., and Cardin, J.A. (2015). Arousal and Locomotion Make Distinct Contributions to Cortical Activity Patterns and Visual Encoding. Neuron 1–15.

Virtanen, P., Gommers, R., Oliphant, T.E., Haberland, M., Reddy, T., Cournapeau, D., Burovski, E., Peterson, P., Weckesser, W., Bright, J., et al. (2020). SciPy 1.0: fundamental algorithms for scientific computing in Python. Nat. Methods 17.

Voigts, J., Sakmann, B., and Celikel, T. (2008). Unsupervised whisker tracking in unrestrained behaving animals. J. Neurophysiol. 100, 504–515.

Voigts, J., Herman, D.H., and Celikel, T. (2015). Tactile object localization by anticipatory whisker motion. J. Neurophysiol. 113, 620–632.

Waiblinger, C., Whitmire, C.J., Sederberg, A., Stanley, G.B., and Schwarz, C. (2018). Primary tactile thalamus spiking reflects cognitive signals. J. Neurosci. 38, 4870–4885.

Wallach, A., Deutsch, D., Oram, T., and Ahissar, E. (2020). Predictive whisker kinematics reveal context-dependent sensorimotor strategies. PLoS Biol. 18, e3000571.

Van Der Walt, S., Colbert, S.C., and Varoquaux, G. (2011). The NumPy array: A structure for efficient numerical computation. Comput. Sci. Eng. 13, 22–30.

Van Der Walt, S., Schönberger, J.L., Nunez-Iglesias, J., Boulogne, F., Warner, J.D., Yager, N., Gouillart, E., and Yu, T. (2014). Scikit-image: Image processing in python. PeerJ 2014, 1–18.

Wang, H.C., LeMessurier, A.M., and Feldman, D.E. (2019). Whisker map organization in somatosensory cortex of awake, behaving mice. BioRxiv 587634.

Wang, Q., Webber, R.M., and Stanley, G.B. (2010). Thalamic synchrony and the adaptive gating of information flow to cortex. Nat. Neurosci. 13, 1534–1543.

Yang, A.E.T., and Hartmann, M.J.Z. (2016). Whisking Kinematics Enables Object Localization in Head-Centered Coordinates Based on Tactile Information from a Single Vibrissa. Front. Behav. Neurosci. 10, 1–15.

Yang, H., Kwon, S.E., Severson, K.S., and O’Connor, D.H. (2016a). Origins of choice-related activity in mouse somatosensory cortex. Nat. Neurosci. 19.

Yang, S.C.-H., Lengyel, M., and Wolpert, D.M. (2016b). Active sensing in the categorization of visual patterns. Elife 5, 1–22.

Yang, S.C.H., Wolpert, D.M., and Lengyel, M. (2016c). Theoretical perspectives on active sensing. Curr. Opin. Behav. Sci. 11, 100–108.

Yau, J.M., Pasupathy, A., Fitzgerald, P.J., Hsiao, S.S., and Connor, C.E. (2009). Analogous intermediate shape coding in vision and touch. Proc. Natl. Acad. Sci. U. S. A. 106, 16457–16462.

Yu, J., Gutnisky, D.A., Hires, S.A., and Svoboda, K. (2016). Layer 4 fast-spiking interneurons filter thalamocortical signals during active somatosensation. Nat. Neurosci. 19, 1–14.

Yu, J., Hu, H., Agmon, A., and Svoboda, K. (2019). Recruitment of GABAergic Interneurons in the Barrel Cortex during Active Tactile Behavior. Neuron 104, 412–427.e4.

Zuo, Y., and Diamond, M.E. (2019a). Rats Generate Vibrissal Sensory Evidence until Boundary Crossing Triggers a Decision. Curr. Biol. 29, 1415–1424.e5.

Zuo, Y., and Diamond, M.E. (2019b). Texture Identification by Bounded Integration of Sensory Cortical Signals. Curr. Biol. 29, 1425–1435.e5.

